# Neurotrophic control of size regulation during axolotl limb regeneration

**DOI:** 10.1101/2021.04.27.441633

**Authors:** Kaylee M. Wells-Enright, Kristina Kelley, Mary Baumel, Warren A. Vieira, Catherine D. McCusker

## Abstract

The mechanisms that regulate the sizing of the regenerating limb in tetrapods such as the Mexican axolotl are unknown. Upon the completion of the developmental stages of regeneration, when the regenerative organ known as the blastema completes patterning and differentiation, the limb regenerate is proportionally small in size. It then undergoes a phase of regeneration that we have called the “tiny-limb” stage, that is defined by rapid growth until the regenerate reaches the proportionally appropriate size. In the current study we have characterized this growth and have found that signaling from the limb nerves is required for its maintenance. Using the regenerative assay known as the Accessory Limb Model, we have found that the size of the limb can be positively and negatively manipulated by nerve abundance. We have additionally developed a new regenerative assay called the Neural Modified-ALM (NM-ALM), which decouples the source of the nerve from the regenerating host environment. Using the NM-ALM we discovered that non-neural extrinsic factors from differently sized host animals do not play a prominent role in determining the size of the regenerating limb. We have also discovered that the regulation of limb size is not autonomously regulated by the limb nerves. Together, these observations show that the limb nerves provide essential and instructive cues to regulate the final size of the regenerating limb.

## Introduction

It is estimated that over 2.1 million Americans are living with limb loss or limb difference, which has profound effects on the function, health, and quality of life in these patients (Ziegler-Graham, MacKenzie, Ephraim, Travison, & Brookmeyer, 2008). The long-term goal of regenerative medicine is to replace or repair damaged limbs by inducing endogenous regenerative responses in humans. Studies in tetrapods capable of regenerating complete and functional limb structures, such as the Mexican axolotl (*Ambystoma mexicanum)*, have been invaluable in terms of understanding the basic underlying biology of limb regeneration and the mechanisms that control this process. Much research in the axolotl system has focused on the essential aspects of the initial stages of regeneration; how the regeneration permissive environment is established, how mature limb cells become regeneration competent, and how the unique pattern of the regenerated limb structures is generated. However, to date, very little is known about the later stages of regeneration that are required for the regenerating structure to mature into a fully functional limb.

One key occurrence at the later stages of limb regeneration is the growth of the limb regenerate to the size that is proportionally appropriate to the rest of the animal. Upon the completion of the developmental stages of regeneration when, the regenerative organ known as the blastema forms, patterns, and differentiates into the missing limb tissues, a proportionally small limb is generated. The regenerated limb then grows rapidly until it reaches a size that is proportionally appropriate to the body length of the animal. Axolotl are an indefinitely growing species, and thus the dimensions of the regenerated limb are different from those at the time of the initial injury. This means that size and proportionality must be regulated throughout regeneration, as opposed to determined at the onset of the process. This is reassuring for the prospect of regenerating limbs on humans, where the difference in the size of the developing compared to the adult limb structure is very large, and the dimensions of the limbs depend greatly on the individual and stage of human maturation (Bogin & Varela-Silva, 2010; Fredriks et al., 2005). However, the mechanisms regulating size and growth during axolotl limb regeneration are largely unknown. Understanding these mechanisms in the axolotl will provide key insight into how a fully functional regenerate can be generated on human patients.

Our limited knowledge of how limb size is regulated is based on historic studies by Harrison and Twitty, who cross-transplanted limb buds from the embryos of differently sized salamander species to determine whether size is instructed by tissue autonomous (intrinsic) or non-autonomous (extrinsic) mechanisms (Harrison, 1924; Twitty & Schwind, 1931). These experiments revealed that both intrinsic and extrinsic factors provide instructive cues to the developing limb bud. More recently, Bryant et al. was able to decouple axolotl limb size from body size through repeated removal of the limb bud (2017). When the limb buds were allowed to regenerate and develop into a limb, they generated permanently miniaturized limbs (D. M. Bryant et al., 2017). Thus, manipulations to the embryonic limb bud can both positively and negatively affect the overall size of the limb on adult salamanders. The factors that regulate tetrapod limb size during embryogenesis, and whether they are reutilized in regenerating limbs, are not known. Correlations between innervation abundance and limb size are apparent in humans and amphibians (Bain et al., 2012; D. M. Bryant et al., 2017; Frykman & Wood, 1978; Mullen, Bryant, Torok, Blumberg, & Gardiner, 1996; Singer, 1978; Tsuge & Ikuta, 1973). In humans, damage to the limb nerves during birth results in impaired limb growth and size, which can be alleviated by surgical repair of the nerves (Bain et al., 2012). Additionally, the presence of neurofibromas in human arms or hands, which increases the abundance of neural signals, results in the formation of proportionally large digits (Tsuge & Ikuta, 1973; Frykman & Wood, 1978). In axolotls, the permanently miniaturized axolotl limbs described above exhibit decreased relative innervation (D. M. Bryant et al., 2017). The role of innervation during the early stages of amphibian limb regeneration has been well established. Nerve signaling is essential for proliferation in the blastema, and loss of innervation at early stages of regeneration results in the complete regenerative failure (Singer, 1978). At the late-bud blastema stage, the loss of nerve signaling results in the formation of complete, yet miniaturized, limb regenerates (Mullen et al., 1996). Together, these observations indicate that the positive relationship between neural signaling and limb size is conserved among tetrapods. However, it remains unknown whether the limb nerves play a supportive or instructive role in growth and sizing of the limb regenerate.

In the current study, we have performed the first characterization of growth in late (post-blastema) staged regenerating limbs of the Mexican axolotl, have identified two post-developmental growth phases, and have shown that both an increase in cell number and cell size contribute to growth of the limb during these phases. Our data shows that innervation is required to maintain late staged growth, and that changes in the nerve abundance are sufficient to manipulate (positively or negatively) the ultimate size of the limb regenerate. We have developed a new regenerative assay that is a derivative of the Accessory Limb Model (ALM), which decouples the nerve source from the host animal, in an assay that we call the Neural-Modified ALM (NM-ALM). Using this assay, we were able to determine that non-neural extrinsic factors do not play an instructive role in size determination during limb regeneration. Last, our findings indicate that the neural regulation of size requires the nerve cell bodies to remain in their endogenous environment, suggesting that upstream cues either from the tissue environment surrounding the nerves or from the Central Nervous System (CNS) are required. Together, our data indicates that limb nerves play an instructive role on the sizing of the amphibian limb regenerate. These observations will be foundational to future work on the identification of the molecular mechanisms that regulate this process.

## Results

### The axolotl limb undergoes at least three stages of growth during regeneration

The majority of limb regeneration research has focused on the early, developmental stages of limb regeneration, and there was little data on the later stages when the limb regenerate grows and matures into a functional structure. Therefore, we started by characterizing the growth of the regenerate until it had completed regeneration. We measured limb and body length to calculate both limb “proportionality” (ratio of limb length to body length) (Supplemental Figure 1) and the growth rate in 10cm sized animals over a period of 140 days, which is when the regenerated limb was no longer significantly different in size relative to the uninjured limbs on the same sized animal, and the growth rate of these limbs was also equivalent. Because axolotls are indeterminate growers, both their limb and body lengths continue to increase in size during the course of regeneration, thus the ratio of limb length to body length was used to normalize for the changes in animal size as time proceeded.

**Figure 1:**
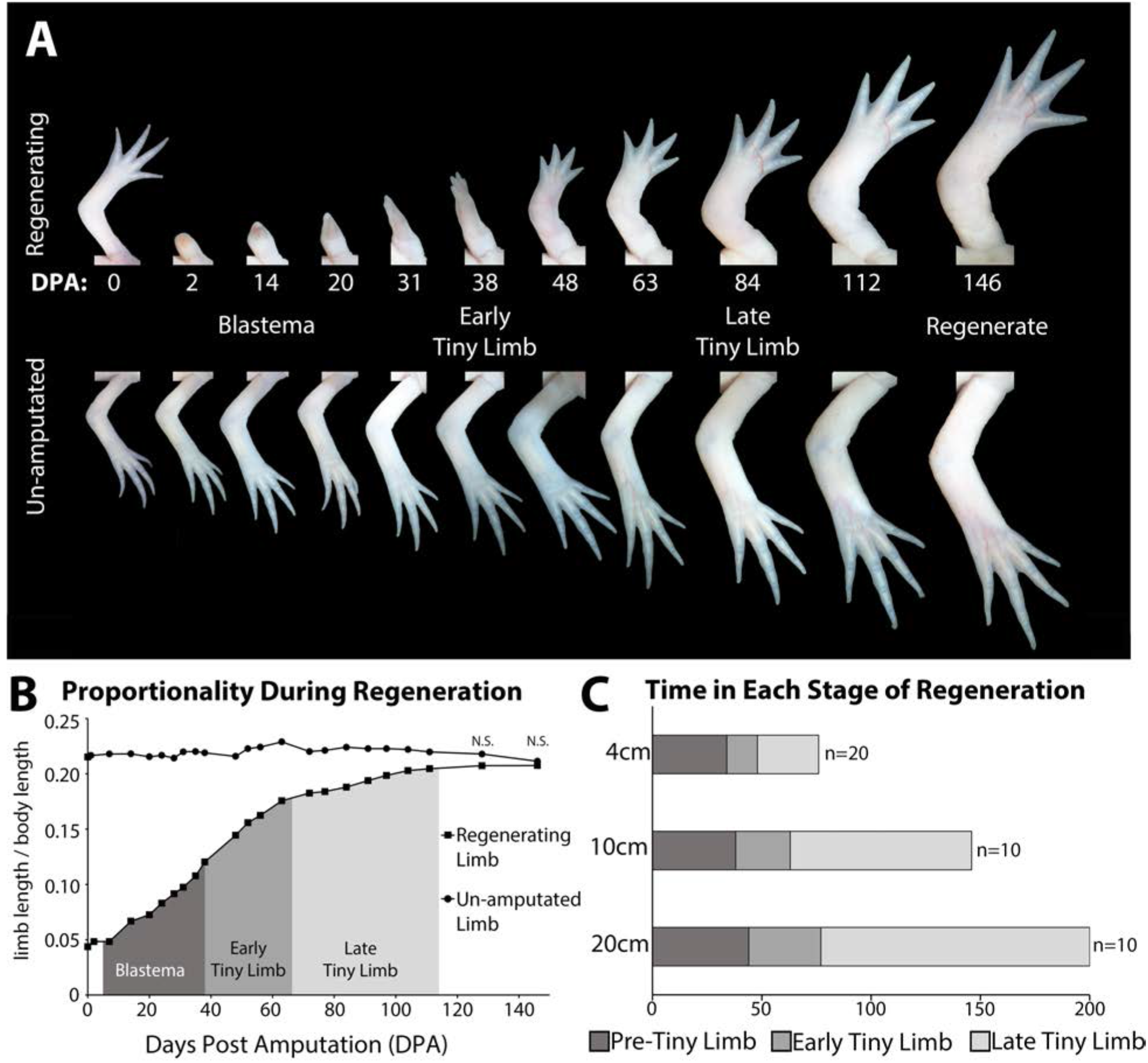
The tiny limb grows at an increased rate compared to an unamputated limb. (A) Time course of growth in amputated (top panel) and the contralateral non-amputated (lower panel) limbs on a 10 cm animal over 146 days. (B) The ratio of limb to body length in regenerating and unamputated limbs was measured over time (10cm animals; n=10). We have separated the growth of the limb regenerate into three stages: the blastema stage (dark grey), the early tiny limb stage (medium grey), and the late tiny limb stage (light grey). Error bars = standard error of the mean. T-Test was used to evaluate significance between the regenerating and uninjured limb size at each time point. All data points <u>not</u> marked with N.S. had p-values less than 0.005. (C) Histogram showing the average amount of time in days that the regenerating limb is in each growth stage for animals of different sizes (4 cm, 10 cm. and 20 cm in length).

We observed that the regenerating limb underwent at least three stages of growth before it reached the size of the unamputated limb: the blastema stage and two post-developmental stages (Figure 1A and B). The blastema stage of growth is comprised of the time immediately following amputation all the way through the digit stage of blastema development. Previous studies have shown that the expression of growth factors and signaling molecules associated with blastema development are lost by the digit staged regenerate, thus indicating the end of the blastema stage, and initiation of post-developmental regenerative processes (Gerber et al., 2018; Nacu, Gromberg, Oliveira, Drechsel, & Tanaka, 2016; A Satoh, Graham, Bryant, & Gardiner, 2008). The digit staged regenerate is significantly smaller than the unamputated limb on sized-matched animals (Figure 1A and B). This small regenerate grew rapidly until it reached the proportionally appropriate size. We have named this post-blastema staged regenerate the “tiny limb.” The tiny limb grows rapidly at a growth rate that is similar to the speed of growth during the blastema stage, 0.04 cm/day (Figure 1B and Supplemental Figure 2). Approximately 4 weeks later (in 10cm sized animals), the growth rate of the tiny limb slows significantly to 0.02 cm/day over the following 9 weeks until both the size and the growth rate of the regenerated limb is not significantly different than the unamputated limb on size-matched animals (Figure 1B and Supplemental Figure 2). Based on this difference in growth rate, we have separated the tiny limb stage of development into two phases of growth: the early tiny limb stage and the late tiny limb stage.

**Figure 2:**
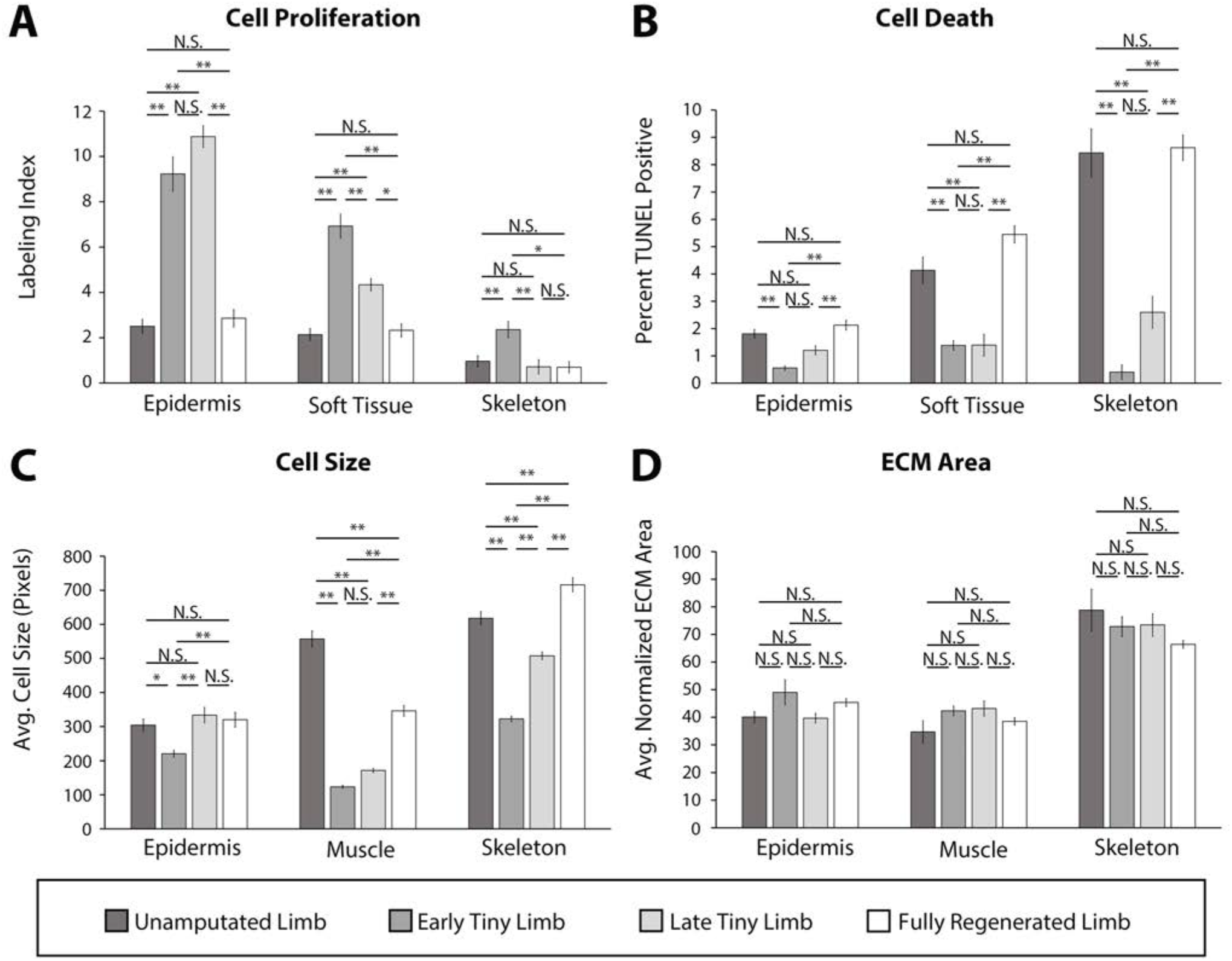
Tiny limb staged regenerates have increased proliferation, decreased cell death, and smaller cell sizes than uninjured or completely regenerated limbs. Transverse sections through the zeugopod of limbs at different stages of regeneration were analyzed for cell proliferation (A), apoptosis (B), cell size (C), and ECM size (D). (A and B) Cell proliferation and death were analyzed in the epidermis, soft tissue, and skeletal elements. A) Cell proliferation was analyzed by EdU labeling (n=5). B) Cell death was analyzed using TUNEL labeling (n=4). (C and D) Cell and ECM size measurements were quantified in the epidermis, muscle, and skeletal elements. C) Cell size was quantified using fluorescently tagged Wheat Germ Agglutinin (plasma membrane) for epidermal and muscular analysis and Alcian Blue staining (collagen) for skeletal analysis (n=4). D) ECM area was calculated by [(tissue area – cellular area)/tissue area] (n=4). Error bars = SEM. P-values calculated by ANOVA and the Tukey Post-hoc test. *=p<0.05 **=p<0.005.

Both the growth rate and the amount of time spent in each of the three stages (blastema, early tiny limb, and late tiny limb) is dependent on the size of the animal at the time of limb amputation. Comparison of these stages between 4, 10, and 20cm animals (snout to tail tip) showed that as body length increases, the growth rate decreases (Supplemental Figure 2) and the amount of time spent in the tiny limb phase increases (Figure 1C). In fact, when comparing animals two-fold different in size, the amount of time in the early tiny limb phase increases by 30-40% (Figure 1C) and growth rate falls by 30-40% (Supplemental Figure 2). This difference in growth rate (measured in terms of regenerate elongation) may be due to the difference in the amount of tissue that needs to be regenerated, since larger animals have more tissue to regenerate than the smaller animals.

### Growth of the tiny limb is mediated by increased cell number and cell size

We next wanted to determine what cellular mechanisms were contributing to growth of the tiny limb. Multiple processes can contribute to tissue growth including increased cell number via regulation of cell proliferation and apoptosis, increased cell size, and extra cellular matrix (ECM) deposition (Conlon & Raff, 1999; Leevers, Weinkove, MacDougall, Hafen, & Waterfield, 1996; Penzo-Mendez & Stanger, 2015; Stanger, 2008; Stocker & Hafen, 2000). Thus, we quantified each of these processes during the different phases of regeneration, relative to uninjured limbs, to determine which could be contributing to growth of the tiny limb. Additionally, we speculated that the contribution of these cell processes could vary in the different tissue types in the regenerating limb. Rather than quantifying the above-described processes globally, we analyzed the epidermis, soft tissue (including all tissues except for skeleton and epidermis), and skeletal tissue (bone and cartilage) separately.

Regenerating limbs at the different stages of growth were sectioned transversely mid-zeugopod, and cell proliferation and cell death were each analyzed by either EdU incorporation or TUNEL staining, respectively. We observed significantly more cell proliferation and significantly less cell death in all tissues analyzed in the early and late tiny limb staged regenerates compared to the unamputated and fully regenerated limbs (Figure 2A and B). Interestingly, the fold increase or decrease in cell proliferation or death, respectively, differed depending on the tissue type. The largest increase in cell proliferation was observed in the epidermis (3.7-fold increase, Figure 2A), while soft tissue and skeletal tissue had slightly more modest increases (3.2 and 2.4-fold increases respectively, Figure 2A). The largest decrease in apoptosis was seen in the skeletal tissue (20.8-fold decrease, Figure 2B), while the soft tissues and epidermis exhibited more moderate decreases of 3.0 and 3.3-fold, respectively (Figure 2B).

Cell size and ECM area were analyzed using a combination of fluorescent and histological stains on sectioned limbs. Wheat Germ Agglutin was used to label the plasma membrane of the epidermal and muscle cells, and Alcian blue stained the ECM of the skeletal tissue (Supplemental Figure 2) (more detail in materials and methods). To standardize our quantification of the average cell size, we measured only the area of cells where the nucleus was observable (Supplemental Figure 2A). We observed that cell size was significantly smaller in the regenerating tissue than the uninjured tissue and increased as regeneration progressed (Figure 2C). This was most profound in the muscle (4.5-fold smaller) and least in the epidermis (1.4-fold smaller, Figure 2C). The ECM area was calculated indirectly by subtracting the total cellular area from the tissue area and dividing by the tissue area (Supplemental Figure 2B) (more detail in materials and methods). However, we did not observe any significant differences in the extracellular compartment of limbs, indicating that ECM deposition does not play a significant role in growth of the tiny limb (Figure 2D).

Together, this data indicates that a combination of increased cell proliferation, decreased apoptosis, and increased cell size contributes to the growth of the tiny limb staged regenerate. Additionally, while all tissue types showed the same trends in all of the cell processes that we analyzed, our data suggests that different cell processes contribute more or less to growth in different tissue types. Future studies will be required to resolve these tissue-specific contributions to growth in more detail.

### Growth of the Tiny Limb is dependent on limb nerves

One interesting observation from the above-described characterization is that the abundance of cell proliferation, cell death, and cell size all show similar trends regardless of the tissue type assessed during each stage of growth in the regenerate. This suggests that there could be a singular signal that coordinates these processes such that the highest growth-promoting signal is occurring during the early tiny limb stage when growth is most abundant and decreases as the growth rate slows during the late tiny limb stage. Thus, we next sought to determine the source of the signal that regulates cell proliferation, death, and size during regeneration.

Previous studies indicate that nerve signaling is required for growth in developing and regenerating limbs. The role and mechanism of neurotropic regulation during the early (blastema) stages of regeneration has been widely studied (Farkas, Freitas, Bryant, Whited, & Monaghan, 2016; Farkas & Monaghan, 2017; Kumar, Nevill, Brockes, & Forge, 2010; Makanae, Mitogawa, & Satoh, 2014; Singer, 1946, 1952, 1978; Singer & Inoue, 1964). It has been well established that nerve signaling is required for, and is a key driver of, blastemal cell proliferation (Brockes, 1984; Brockes & Kintner, 1986; Lehrberg & Gardiner, 2015). Furthermore, generation of permanently miniaturized limbs through repeated removal of limb buds results in “mini limbs” that are hypo-innervated compared to controls (D. M. Bryant et al., 2017). We therefore hypothesized that nerves could play a role in driving growth during the tiny limb stages of regeneration.

To test this idea, we first characterized the abundance of innervation during these post-developmental stages of regeneration. We collected early and late tiny limbs, as well as unamputated and fully regenerated limbs for comparison, and sectioned them transversally through the zeugopod. The sections were stained with an anti-acetylated tubulin antibody (nerve – green, Figure 3A), and we measured the abundance of innervation relative to the limb area (Figure 3B). This quantification revealed significantly higher levels of relative innervation during the early and late tiny limb stages compared to unamputated and fully regenerated limbs (Figure 3B). Interestingly, the total abundance (not normalized to tissue area) of innervation increases from the early to late tiny limb stages (Supplemental Figure 4). Thus, the decrease in relative innervation during the transition from the early to late tiny limb stages is likely due to the substantial increase in limb area of the later staged regenerate. The relative abundance of innervation also correlates well with the growth rate in these tissues. The early tiny limb stage has the highest relative abundance of innervation, followed by the late tiny limb stage (Figure 3B). The abundance of innervation of the completed regenerate has decreased to that of the uninjured limb (Figure 3B). We speculated that nerves could provide growth-promoting signals during regeneration, which decreases as the relative abundance of innervation decreases, slowing the growth of the regenerate as it reaches its final size.

**Figure 3:**
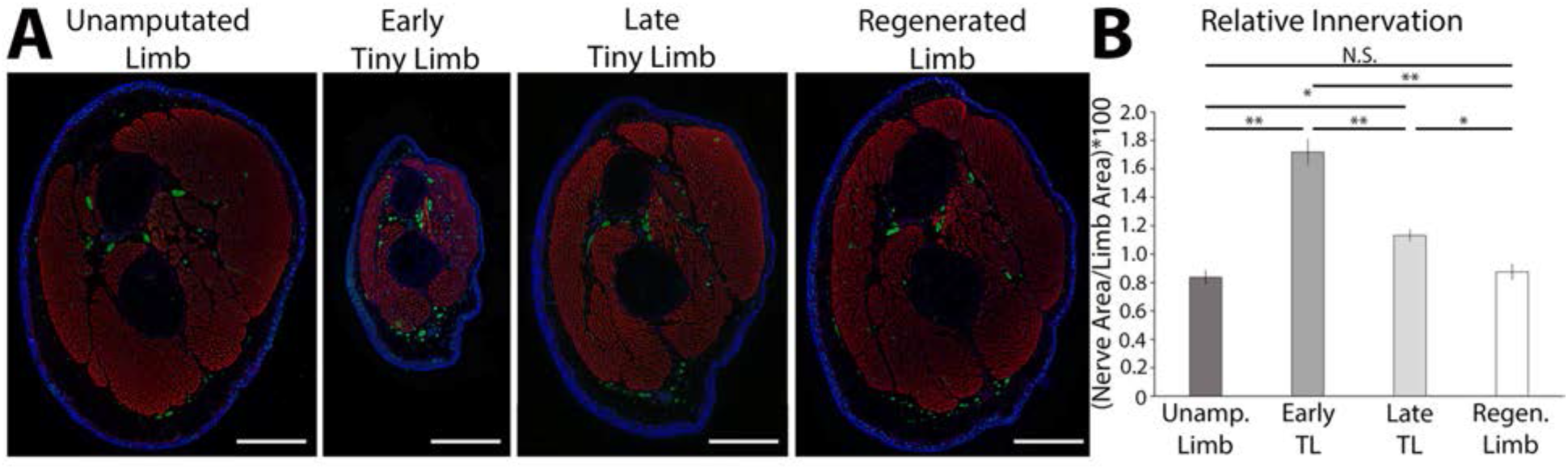
The tiny limb staged regenerate is hyperinnervated. A) Fluorescent images were obtained of transverse sections of uninjured, early and late tiny limb stages, and fully regenerated limbs (DAPI = blue, Phalloidin (for actin filaments) = red, Acetylated-tubulin (for nerves) = green; scale bars are 1000um). B) Nerve area relative to total limb area was quantified from the sections represented in A (n=5). Error bars = SEM. P-values calculated by ANOVA and the Tukey Post-hoc test. *=p<0.05 **=p<0.005.

**Figure 4:**
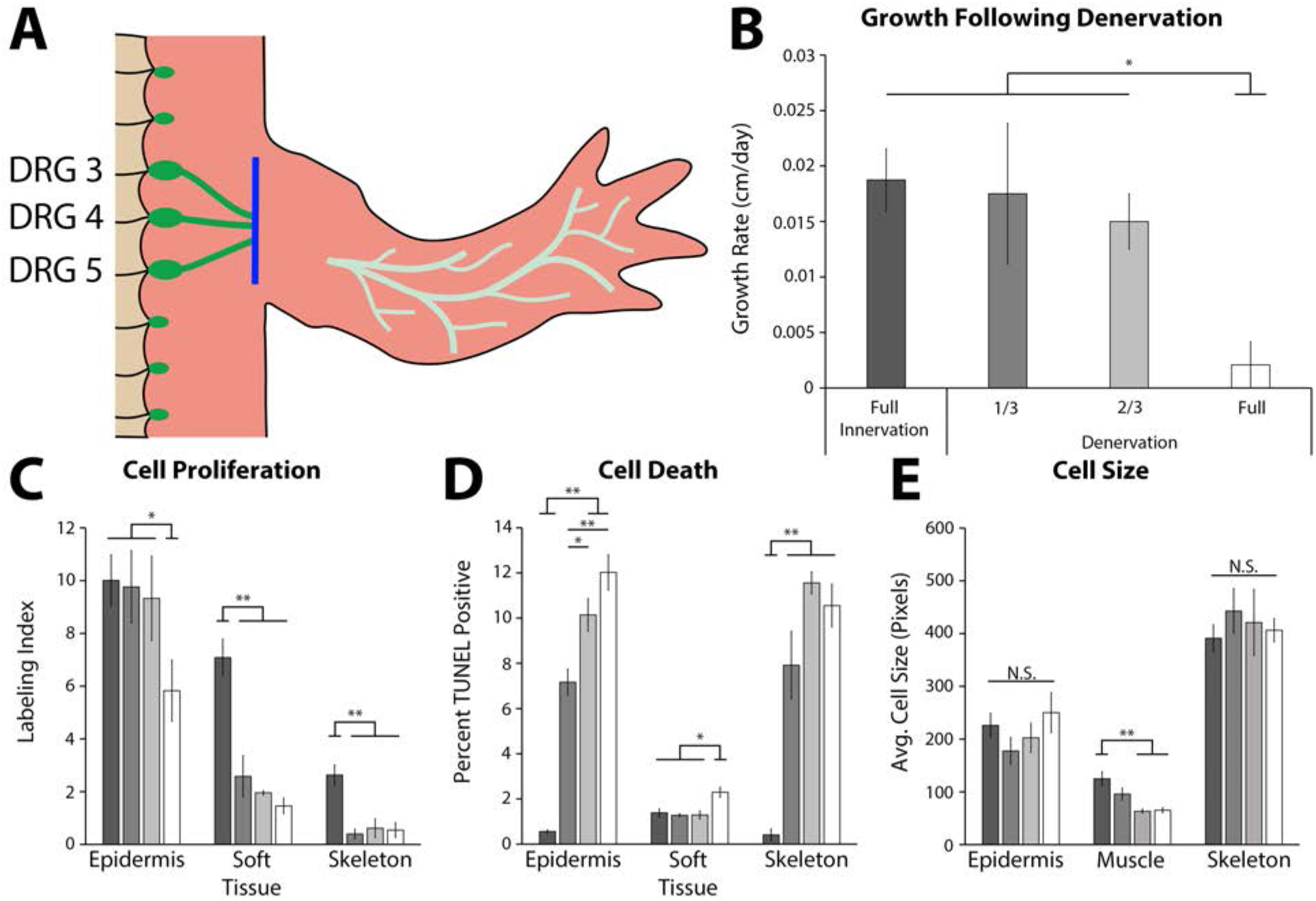
Innervation is required for growth of the tiny limb staged regenerate. A) Dorsal root ganglia (DRGs) 3, 4, and 5 (green dots) are located lateral to the spinal column and their nerve bundles (green lines) feed into the forelimbs. Limbs were amputated and permitted to regenerate to the early tiny limb stage, at which point, either a mock, partial (1/3 = DRG 5 or 2/3 = DRGs 4 and 5), or full denervation (represented) was performed by severing (blue line) and removing sections of the nerve bundles. Limbs were collected 4 days post denervation, and growth rate (B), cell proliferation (C), cell death (D), and cell size (E) were analyzed for limbs with mock denervations (n=6), 1/3 denervations (n=5), 2/3 denervations (n=5), and full denervations (n=6). The color of the bars in panels C-E refers to the color of the bars in panel B. Error bars = SEM. P-values calculated by ANOVA and the Tukey Post-hoc test. *=p<0.05 **=p<0.005.

To determine whether nerve signaling plays a functional role in determining size, we next tested whether nerve signaling is required to maintain growth in the tiny limb staged regenerate. Limbs were amputated and permitted to regenerate to the early tiny limb stage, at which point nerve signaling was severed (blue line, Figure 4A) via denervation at the brachial plexus. To test for a possible dose response or signaling threshold effect, we severed either one, two, or all three of the nerve bundles at the plexus. Mock denervation surgeries were performed as controls. The limbs were measured prior to denervation and four days post denervation, when they were collected for analysis. The growth rate, abundance of cell proliferation, abundance of cell death, and cell size were all analyzed (Figure 4B-E).

We observed that nerve signaling is required for growth of the tiny limb. Fully denervated early tiny limbs had a 9-fold slower growth rate than innervated early tiny limbs (Figure 4B). Likewise, cell proliferation was negatively impacted by denervation in all tissue types analyzed (1.7, 4.9, and 4.8-fold decreases in the epidermis, soft tissue, and skeleton respectively; Figure 4C). Cell death levels in all tissues were significantly increased following full denervation (21.6, 1.6, and 25.7-fold increases in epidermis, soft tissue, and skeleton respectively; Figure 4D). Lastly, cell size appears to only be significantly affected in the muscle, where there is a near 2-fold decrease in average cell size in the denervated early tiny limbs (Figure 4E). When late tiny limbs were fully denervated, only the growth rate and abundance of cell proliferation were significantly decreased (Supplemental Figure 5). Thus, neurotrophic regulation of cell death and cell size (in the muscle tissue) appears to be restricted to the early tiny limb stage of growth.

**Figure 5:**
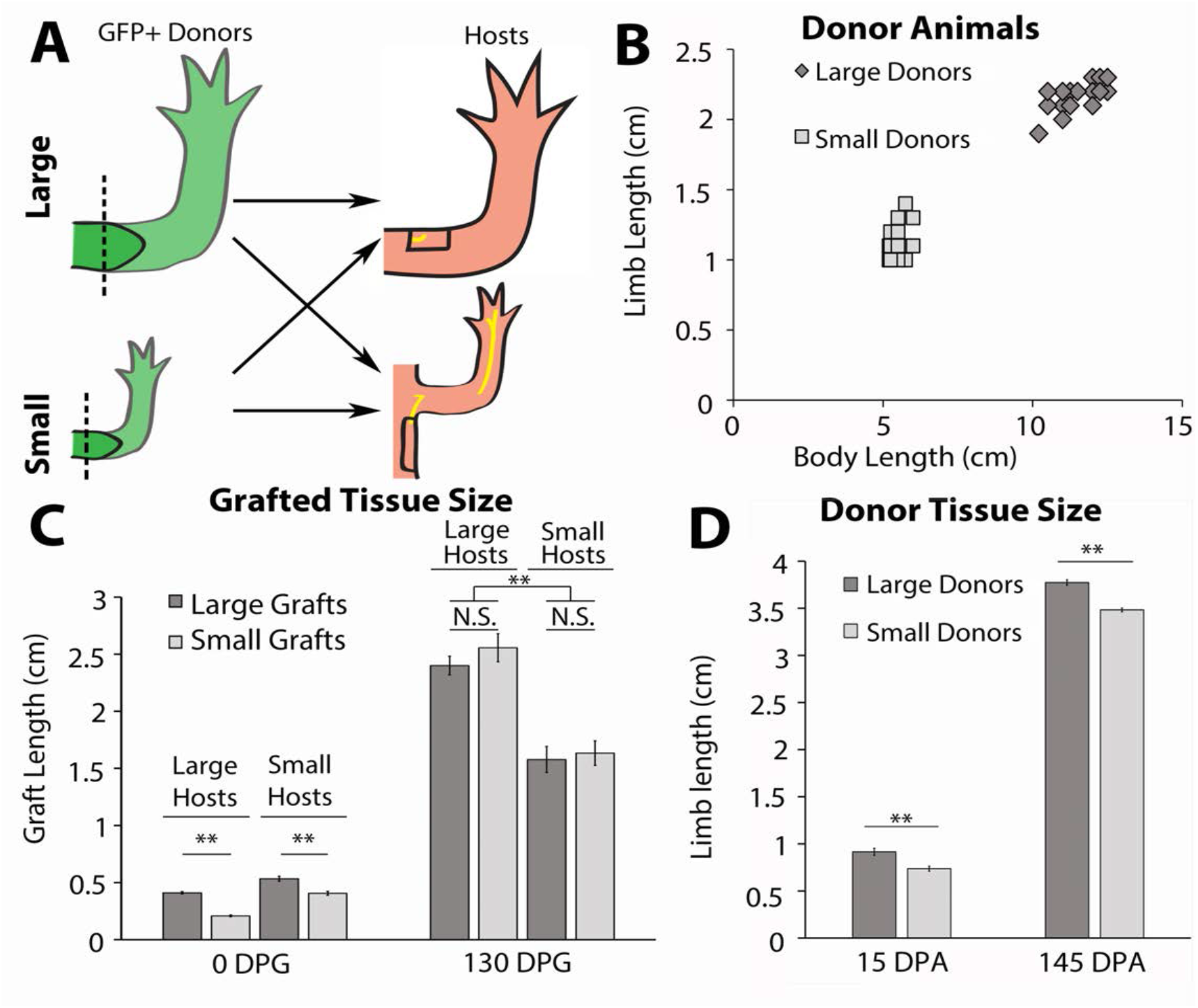
Innervation abundance determined regenerate size. A) Blastemas with approximately 2mm of stump tissue from large and small GFP+ donor animals were grafted onto a regenerative permissive environment, a wound site with a deviated limb nerve bundle, on large or small host animals. B) Limb length and body length were measured on the GFP+ donor animals. C) The regenerating grafted tissues were measured at 0 and 130 DPG. Blastemas from large donors (dark grey) were grafted onto large (n=7) and small (n=9) host animals, and blastemas from small donors (light grey) were grafted onto large (n=9) and small (n=10) host animals. D) The regenerating large (n=10) and small (n=20) animal donor limbs were measured at 15 and 145 days post amputation. Error bars=SEM. P-values calculated by ANOVA and the Tukey Post-hoc test. *=p<0.05 **=p<0.005.

The partial denervations revealed either a dose response or threshold response depending on the tissue and growth characteristic quantified. A dose response is reflected by a linear relationship between abundance of signal and the phenotype. A threshold response indicates a specific abundance of a signal is required for a phenotype, for example, a specific level of innervation is required for growth. We observed that cell proliferation in the soft tissue and skeleton decreased significantly, to full denervation levels, with partial denervations indicating that there is a high threshold of nerve signaling responsible for maintaining cell proliferation in these tissues (Figure 4C). Conversely, cell death in the epidermis had a strong dose response, with significant incremental increases with increased denervation (Figure 4D). These results indicate that each tissue responds differently to nerve signaling to maintain growth. This could explain the decreasing growth rate trend in partial denervations, which only becomes significant with the full denervation (Figure 4B). Together these results reveal the complexity of neuronal regulation of growth and indicate that an evaluation of the tissue-specific responses to nerve signaling is required for a complete understanding of growth and size regulation during regeneration.

### Innervation abundance determines regenerate length

Having established that nerve signaling is required to maintain growth during limb regeneration, we next wanted to determine whether we could positively and negatively manipulate the size of the regenerate by altering the abundance of innervation. To test this, we performed a grafting experiment between differently sized axolotl (Figure 5A). Limb nerve bundles increase in size as the animal grows, and quantification of the cross-sectional area of the nerve bundles extending out from the limb Dorsal Root Ganglia (DRGs) reveals that the size of the bundle is almost 2-fold larger in 14cm long animals compared to 7cm animals (Supplemental Figure 6). Thus, blastema grafts onto the limbs of large hosts will be innervated by larger nerve bundles than grafts on small host animals.

**Figure 6:**
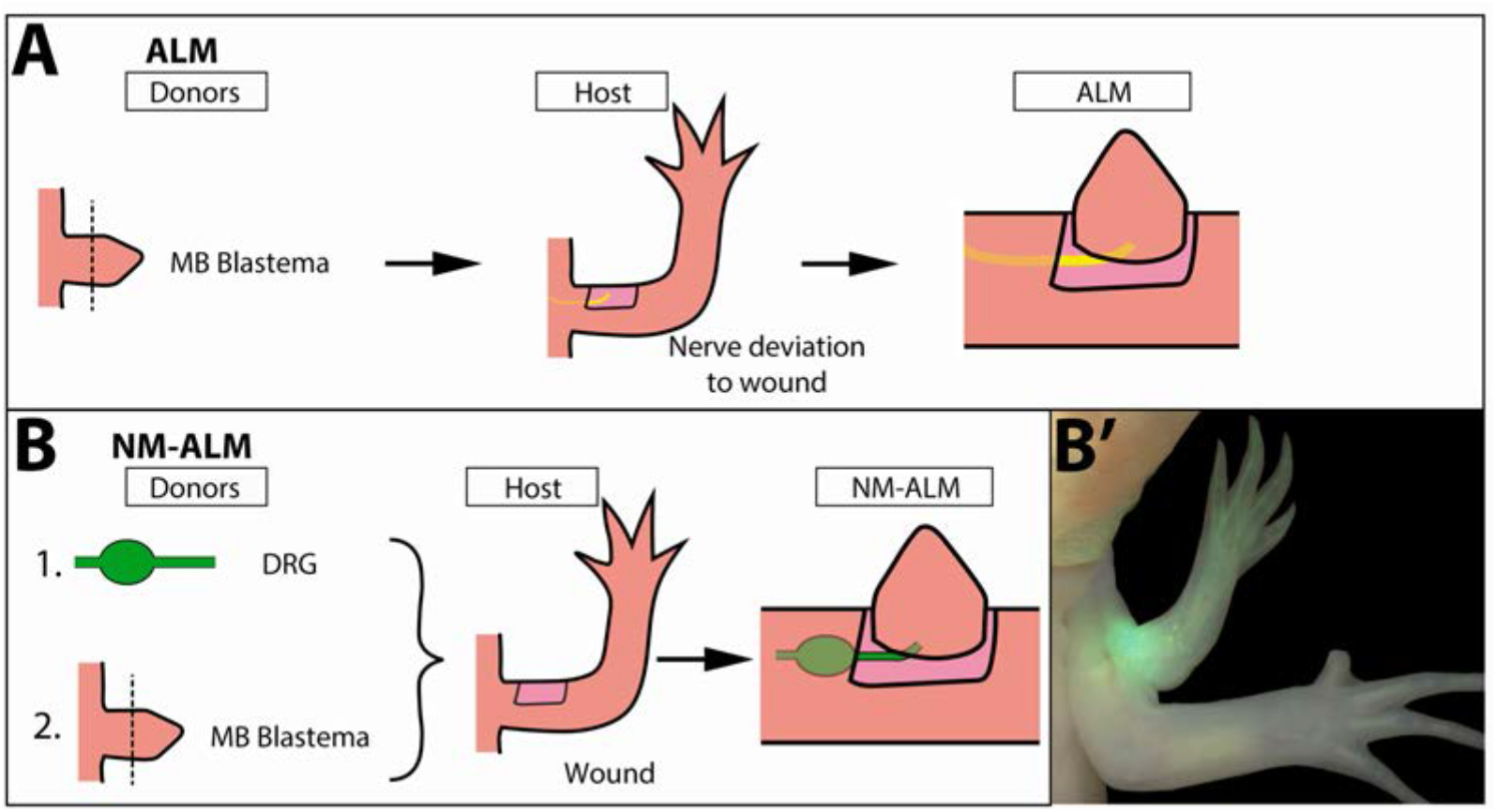
Decoupling host environment with innervation using the Neural-Modified ALM (NM-ALM). A) The traditional ALM (as used in Figure 5) requires a blastema donor and a host animal with a nerve bundle deviated to the wound site. The NM-ALM requires a GFP+ DRG donor, blastema donor, and host limb with a wound site. B’) The DRG’s GFP+ axons regenerate and innervate the ectopic limb.

To generate large and small animals we housed age-matched GFP+ (donors) and GFP-(hosts) axolotl at either 19°C or 4°C. Housing half the animals at 4°C stunted their growth while the 19°C animals grew at a faster rate. After approximately two months, the 19°C animals were 2-fold larger in body length and limb length than the 4°C animals (Figure 5B). After the large and small siblings were both incubated at 19°C for 14 days, both forelimbs on small (~6cm) and large (~12cm) GFP+ axolotl were amputated and permitted to regenerate to the mid-bud blastema stage (Figure 5A and B). The blastemas and approximately 2mm of stump tissue were grafted onto regenerative permissive environments on small (~6cm) and large (~12cm) GFP-host animals (Figure 5A). It has previously been found that cells from approximately 500μm of stump tissue migrate and contribute to the regenerate (Currie et al., 2016). Therefore, stump tissue was included with the blastemas to prevent the contribution of cells from the host environment to the regenerate.

A regeneration permissive environment on the large animal hosts was generated by deviating the branchial nerve bundle to an anterior located wound site on the limb, using the standard Accessory Limb Model surgery (Figure 5A) (Endo, Bryant, & Gardiner, 2004; McCusker & Gardiner, 2013). Because the limb circumference of the small host animals was too small to receive the grafted blastema tissue from the large donors, we deviated the branchial nerve to a wound site on the flank of the host, where a larger skin wound could be made to fit the larger graft size (Figure 5A). The length of the ectopic limbs was measured bi- or tri-weekly. They were considered fully regenerated when the ectopic limb growth rate was no longer significantly different from the limbs of the host animals. The donor animal limbs that were amputated for the blastema grafts were also continually measured during regeneration as an additional control to determine limb sizes of the large and small blastemas if left on their native environment.

We hypothesized that if nerve abundance can regulate size of the limb regenerate, we would observe that the lengths of the grafted regenerates will correspond to the size of the host environment rather than the donor. Thus, blastemas from the small donors will produce large ectopic limbs when grafted to large hosts, and blastemas from large donors will produce small ectopic limbs when grafted to small hosts. Alternately, if nerve abundance does not influence regenerate size, then we would expect to see the blastemas from small donors on large hosts produce ectopic limbs smaller than the control grafts from large animals, and vice versa.

We observed that nerve abundance influenced ectopic limb size. The blastemas (15 days post amputation) from small donors were initially 2-fold smaller than those from large donors when they were grafted onto the host environments (Figure 5C, left panels, 0 days post graft (DPG)). Interestingly, we observed that the grafted tissues were the same size by 18 DPG within each host type (Supplemental Figure 7A). By approximately 38 DPG, there was a significant difference in size between the ectopic limbs on the large and small host animals, and this continued until 130 DPG, when the growth rate of the grafted limbs matched that of the host limbs (Figure 5C, right panels; Supplemental Figure 7A). In comparison, the control regenerates on the donor animals remained significantly different in size throughout regeneration (Figure 5D). These data indicate that the sizing of the limb regenerate positively correlates with nerve abundance. However, it does not rule out other potential influences from the host environments that may also contribute to size. Thus, we next designed an experiment that decouples nerve abundance from the size of the host to test whether non-neural signals contribute to the sizing of the limb regenerate.

**Figure 7:**
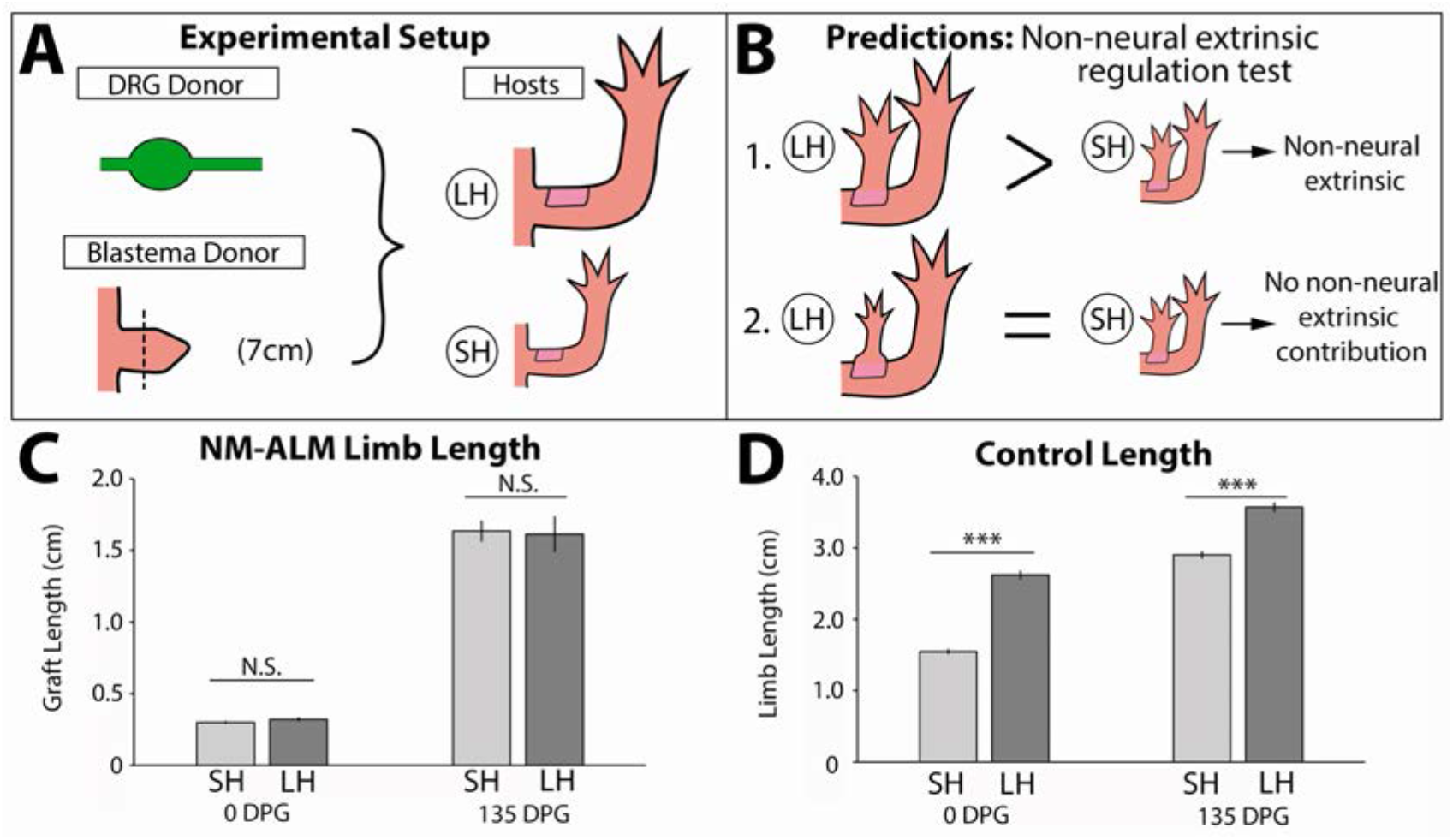
Non-neural extrinsic factors do not play an instructive role in size regulation. A) DRGs from GFP+ donor animals (~14cm) were grafted into wound sites on large (~14cm, n=12) and small (~7cm, n=19) host animals followed by mid-bud blastemas from (~7cm) donor animals. B) If non-neural extrinsic factors play an instructive role in size regulation, then the large host animals would produce a larger ectopic limb than those on the small host animal. If non-neural factors do not play an instructive role then the ectopic limbs will be the same size, regardless of host size. C) The ectopic limb lengths were the same size at 0 DPG and remained the same throughout regeneration (135 DPG). D) The unamputated limbs on the control animals remained different sizes throughout the course of the experiment (n=10). Error bars=SEM. P-values calculated by T-tests. ***=p<0.0005.

### Non-neural extrinsic signals do not provide instructive cues on regenerate size

To evaluate the potential role of non-neural sources of size regulation that may be present in the differently sized host animals we developed a new regenerative assay called the Neural Modified Accessory Limb Model (NM-ALM) that decouples the source of the nerves from the host environment (Figure 6B). In the NM-ALM, the limb Dorsal Root

Ganglia (DRGs) from GFP+ donor animals are harvested and grafted into lateral limb wounds on host animals (one DRG per wound). The GFP+ DRGs are grafted below the mature skin next to the wound site, and the axon bundles are pulled into the middle of the wound site in a similar manner as the traditional ALM surgery. Mid-bud blastema staged regenerates from age-matched donors are then grafted onto the wound site (Figure 6B). We measured the length of the ectopic limbs in the NM-ALM weekly or bi-weekly, and their growth rates (cm/day) were compared to the unamputated limbs on the donors. The grafted limbs were considered to have completed limb regeneration when their growth rates were not significantly different from the growth rates of the unamputated limbs on the control animals. We observed that the implanted DRG supported the continued development of the blastema into a completely patterned and differentiated limb (Figure 6B’). Thus, the implanted DRGs are able to provide the appropriate signals to support the regenerative process.

To test whether non-neural extrinsic signals provide instructive cues that regulate the size of the regenerating limb we performed the NM-ALM on differently sized host animals (Figure 7A). If non-neural extrinsic signals play a role in regulating the size of the regenerate, then we expected to observe that the regenerates that grew in the NM-ALMs on the large (14 cm) hosts would be larger in size than the ones that grew on the small (7 cm) hosts (Figure 7B). Alternately, if signaling from the nerves play the major instructive role then we would not expect to see differences in the size of the regenerates on the different sized hosts. After 135 days post grafting, we observed that there continued to be no significant difference in the length of the grafted regenerates on the different sized host animals (Figure 7C). This trend was observed in NM-ALMs that were performed with DRGs that were harvested from both large and small animals (Supplemental Figure 7B). In contrast, the control uninjured limbs on the small and large animals remained significantly different in size (Figure 7D). Because the size of the limb regenerate was not impacted by the host environment when the abundance of innervation was constant, we conclude that non-neural extrinsic factors play a negligible role on instructing the size of the regenerate. We, however, are not ruling out the possible (and likely) supportive role that other extrinsic factors may play in the growth of the regenerate.

### Size regulation cues from limb nerves are not autonomous

To test whether the neural regulation of regenerate size is autonomous or not, we leveraged our newly developed NM-ALM assay using DRGs that were harvested from different sized GFP+ donors (14cm versus 7cm) (Figure 8A). Our expectation was that if neural regulation of size occurs autonomously (at the DRG-level), then the size of the regenerated limb that grew on the DRG from the large donor would be larger than that which grew on the one from the small donor (Figure 8B). Conversely, if the DRGs require a connection with their native environment to regulate regenerate size, then we expected to see no difference in the regenerate size regardless of the donor source of the DRG (large or small) (Figure 8B). Because we had previously observed a dose response in limb growth in partially denervated regenerating limbs (Figure 4), we hypothesized that limb size could be regulated by nerve abundance alone. However, our data showed that there was no statistical difference in ectopic limb length between the regenerates generated from NM-ALMs implanted with the DRGs from small or large animals (Figure 8C). This trend was observed on the NM-ALMs on both the large and small hosts (Supplemental Figure 7B). Thus, we concluded that factors from the DRGs’ endogenous environment are required for the neural regulation of regenerate size. Future studies will focus on the cells and signals that play a role in this upstream regulation.

**Figure 8:**
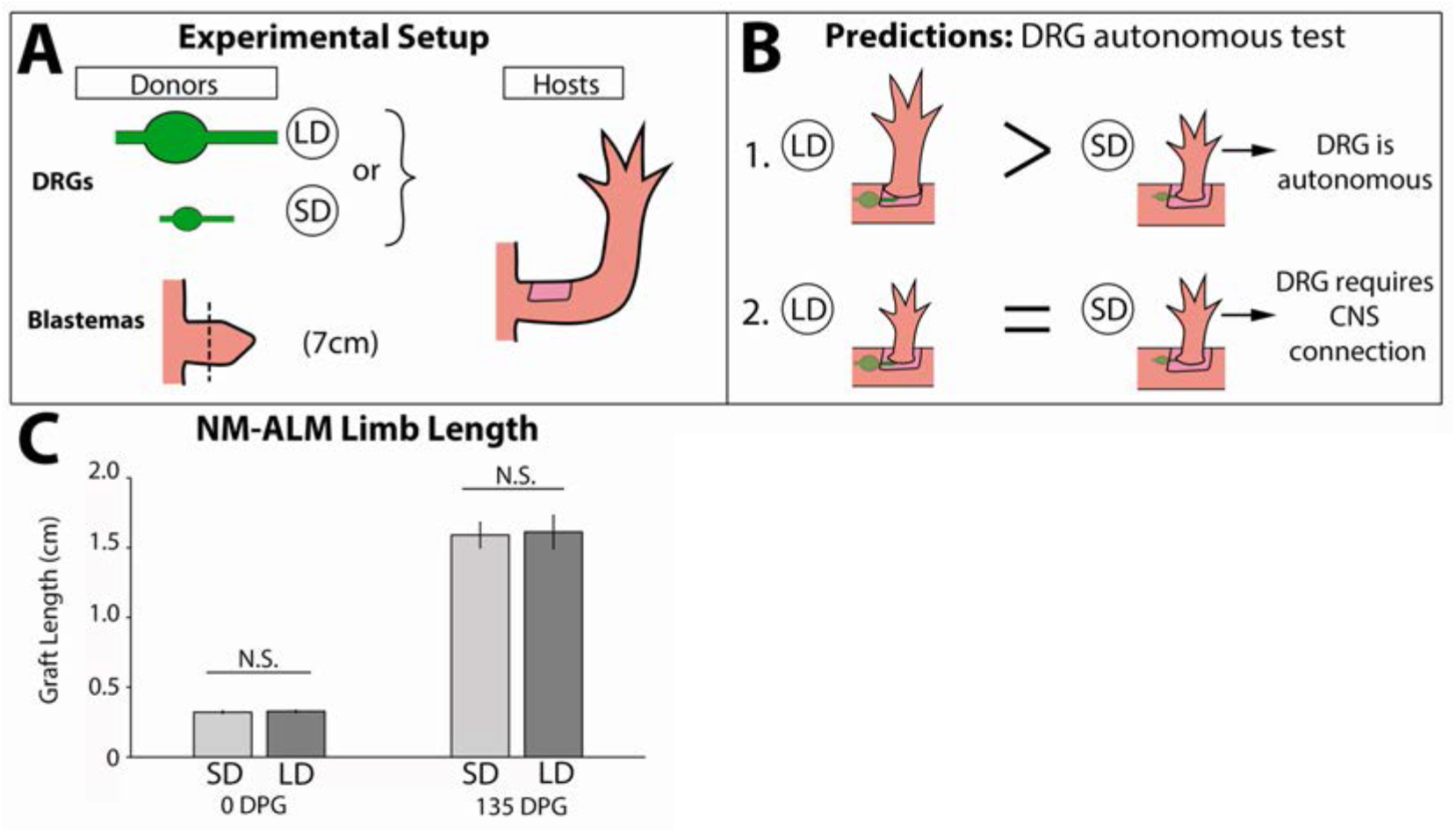
Neural-regulation of regenerate size is lost upon separation from spinal column. A) DRGs from large (~14cm, n=12) and small (~7cm, n=16) GFP+ animals were grafted into wound sites on host animals(~14cm) followed by mid-bud blastemas from (~7cm) donor animals. B) If the ability to regulate size is autonomous to the DRGs, the DRGs from large animals will produce larger ectopic limbs than those from small animals. If size regulation is not autonomous, there will be no difference in ectopic limb size between grafts supplied by large or small animal DRGs. C) The ectopic limb lengths were the same size at 0DPG and remained the same throughout regeneration (135DPG). Error bars=SEM. P-values calculated by T-tests.

## Discussion

One of the last steps of regeneration required to generate a fully functional limb is the growth of the regenerate to a size that is proportionally appropriate to the animal. However, little data had previously been collected on the post-blastema stages of regeneration and how growth and size are regulated. This study constitutes the first thorough investigation of how the regenerating limb grows to the proportionally appropriate size. We have discovered that there are three distinct phases of growth prior to the limb completing regeneration; the blastema phase, the early tiny limb phase, and the late tiny limb phase (Figure 1). During the tiny limb phases of growth, the regenerate grows through increased cell proliferation, survival and cell size (Figure 2), and innervation is required to maintain this growth (Figure 3 and 4). Our data indicates that nerves play the instructive role in size regulation (Figure 5 and 7), and that factors from the native neural environment are required for this regulation of regenerate size.

### Size determination during limb development versus limb regeneration

Previous studies on developing amphibian limbs suggest that both intrinsic and extrinsic factors determine size during embryogenesis. One example is from the classic cross-grafting study performed by Ross Harrison using limb buds between two differently sized salamander species (Harrison, 1924). In this experiment, limb buds from *Ambystoma tigrinum* (adult body length of 27cm) were grafted to *Ambystoma punctatum* (adult body length of 16 cm) hosts and vice versa (Harrison, 1924). It was observed that the grafted limb buds grew to sizes that failed to correspond with limbs from either the donor or the host. Rather, the limb buds from *Ambystoma punctatum* grew to a size that was smaller than the both the host and donor limbs when grafted to *Ambystoma tigrinum*, while the *Ambystoma tigrinum* limb buds grew larger than the host and donor limbs when grafted to *Ambystoma punctatum*. These observations indicate that limb size is determined through both the intrinsic growth capacity of the cells and an extrinsic growth promoting factor that is present in different abundances in the embryos of different host species (Harrison, 1924). It was later shown that when the host animals were fed to capacity, the growth rates of the grafted limbs mimicked the donor animal’s growth rate, but final size of the limb was more closely matched to the host (Twitty & Schwind, 1931). Thus, these classic studies demonstrate that limb size can be influenced by environment, such as available nutrients, during embryonic development.

While our study clearly shows that extrinsic signals from the adult limb nerves play a central role in determining size of the regenerate, it is unknown whether the same nerve-dependent mechanisms are at play during limb bud development in the axolotl. During embryonic limb development, the Apical Ectodermal Ridge (AER) signaling center produces factors such as FGFs and BMPs, which maintain an undifferentiated state and drive proliferation in the limb bud mesenchyme. The AER is established in an aneurogenic environment in *Xenopus laevis, Rana pipiens, Ambystoma punctatum*, and *Ambystoma maculatum* (Keenan & Beck, 2016; Kumar et al., 2011; Nieuwkoop & Faber, 1958; Taylor, 1943; Yntema, 1959), while regenerating limbs require signaling from the nerve to establish the analogous structure, the Apical Epithelial Cap (AEC) (Singer, 1978; Singer & Inoue, 1964). However, axolotl develop limb buds as larvae, not as embryos, and it is unknown whether the axolotl limb bud is innervated or not. Thus, it is possible that the regulation of limb size in the axolotl limb bud and blastema are both dependent on nerve signaling. Regardless, it remains unknown whether the same molecular signals are regulating size in both embryonic and regenerating limbs. As more information on the molecular mechanisms of this process are pieced together, comparative studies can determine conservation between development and regeneration in different amphibian species.

It should be noted that our study does not rule out the influence of intrinsic factors that regulate the size of the regenerating limb. Because our study was performed on sibling *Ambystoma mexicanum*, there were minimal genetic differences between donor and host animals. Performing a similar cross grafting experiment, as was performed by Ross Harrison (1924), between species of different sizes in the context of regeneration would yield valuable insight in this regard.

### Potential role of other extrinsic factors in determining size of the limb regenerate

Our studies indicate that, as opposed to the limb nerves, other extrinsic factors do not play an instructive role in size regulation (Figure 7). However, we have not ruled out the potential contributions of non-neuronal extrinsic factors in supporting the growth of the regenerate. These could include bioelectric signals (Blackiston, McLaughlin, & Levin, 2009; Levin, 2012), hormones (Penzo-Mendez & Stanger, 2015), and circulating growth factors (P. J. Bryant & Simpson, 1984; Conlon & Raff, 1999). For example, bioelectric signals are key regulators of size and proportionality in zebrafish fins, where mutations that generate a leaky potassium channel results in an overgrowth phenotypes (Perathoner et al., 2014). Furthermore, regeneration in a short fin phenotype zebrafish line results in longer fins when treated with a Calcineurin inhibitor, affecting function of the same potassium channel (Daane et al., 2018). In these studies, the bioelectric signals are altered autonomously in the tissue; however, bioelectricity has the potential to be regulated systemically through electromagnetic fields and paracrine signaling (Levin, 2012).

It is also possible that systemic growth factors or hormones could support growth during regeneration. For example, humans and other mammals can undergo a process known as catch-up growth, which occurs in pre-adult humans that exhibit stunted growth as a result of exposure to a stressor such as sickness, emotional stress, or malnourishment (Boersma & Wit, 1997; Williams, 1981). Upon the removal of the source of stress the animal undergoes rapid whole-body growth to “catch-up” to the size that they would have been at their stage of development. Catch-up growth is dependent on the increased expression of growth promoting hormones, such as growth hormone (GH) and insulin growth factor (IGF) in the recovering animals, which could be regulated by the nervous system (Boersma & Wit, 1997; Penzo-Mendez & Stanger, 2015). Although there are similarities, there are multiple facets that distinguish the process of limb regeneration from catch-up growth. First, catch-up growth occurs to the entire body, while the growth of the limb regenerate is isolated to the regenerating structure. Additionally, the success of catch-up growth is largely dependent on the duration of the source of stress and the life-stage of the human when the source is removed. In contrast, axolotl are capable of regenerating complete limbs to the appropriate size irrespective of life stage (Figure 1 and Supplemental Figure 2). Regardless, the potential role of neuroendocrine regulation on growth has not yet been tested in regenerating axolotl limbs.

Systemic factors also play a role in compensatory growth mechanisms. Compensatory growth typically refers to re-growth of missing tissue or a particular organ rather than the entire organism (Boersma & Wit, 1997; Holder, 1981). For example, following a partial hepatectomy in mammals, the remaining liver tissue will regenerate to reestablish proper liver volume (Michalopoulos, 2007). The drivers of compensatory growth are often both local and systemic cues that stimulate regrowth through a feedback mechanism until the appropriate size is reached (Boersma & Wit, 1997; Holder, 1981; Williams, 1981). However, there are key differences between limb regeneration and compensatory growth. Compensatory growth involves reactivation of the cell cycle in differentiated cells. Regeneration involves dedifferentiation of mature cells into blastema cells, proliferation, and redifferentiation into the missing tissue (McCusker, Bryant, & Gardiner, 2015). Furthermore, compensatory growth is thought to cease once the original size is restored, whereas limb regeneration proceeds until proper proportionality is reestablished rather than when original size is reached. Therefore, while regeneration is not considered compensatory growth it is possible that similar feedback between the regenerating limbs and circulating factors in the host body play a role in regulating limb size.

### Mechanism of neurotrophic regulation of size during limb regeneration

Observations from multiple species indicate that neural regulation of limb size is a conserved mechanism. Most notably in humans, damage to the limb nerves corelates to decreased limb size, while overabundance of innervation corresponds to limb or digit enlargement (Tsuge & Ikuta, 1973; Bain et al., 2012; Cerrato et al., 2013; Labow, Pike, & Upton, 2016). However, one of the main unanswered questions remaining is how nerves are regulating size specifically during limb regeneration. While the molecular mechanism by which nerves regulate size has not been determined, the role of innervation during the early stages of limb regeneration has been extensively studied and might be drawn upon to provide insight into their role in size regulation. During the early stages of limb regeneration, it has been observed that innervation of the wound epithelium is essential to establish the apical epithelial cap (AEC). The AEC then produces growth factors essential to induce and maintain the cells in a dedifferentiated state and drive blastema cell proliferation (McCusker et al., 2015). Indeed, Satoh *et al*. demonstrated that FGF8 and BMP7 are directly produced by the nerves in the Dorsal Root Ganglia (DRGs) and migrate to the wound epithelium (Satoh, Makanae, Nishimoto, & Mitogawa, 2016). Furthermore, the requirement of nerves for limb regeneration can be supplemented for by cocktails of growth factor proteins including Fibroblast Growth Factors (FGFs) and Bone Morphogenic Proteins (BMPs) (Makanae et al., 2014; Akira Satoh et al., 2016; Vieira et al., 2019), Neuregulin-1 (Farkas et al., 2016), or Anterior Gradient (Kumar, Godwin, Gates, Garza-Garcia, & Brockes, 2007). Although the AEC is no longer present during the post-blastema stages of growth, nerves could both generate and induce the expression of similar growth promoting factors in the late-staged regenerating tissues. Future studies will focus on identifying the molecular mechanism of neurotrophic regulation of size during limb regeneration.

### Conclusion

This study provides foundational knowledge on the later stages of limb regeneration to understand the how size and proportionality becomes reestablished in a continually growing system. Our data indicates that nerve signaling plays an instructive role in determining regenerate size, and future studies will focus on identifying the molecular mechanism of size regulation. Furthermore, our data suggests that there is an upstream driver of size regulation, potentially in the DRG’s endogenous environment or from the CNS. Lastly, as previously stated, our studies have not ruled out the likely intrinsic factors involved in size regulation. In total, there are still many unknowns, both up and downstream of nerve signaling, that must be resolved to fully understand how size and proportionality become reestablished during axolotl limb regeneration. Furthermore, as regenerative medicine seeks to tap back into the developmental mechanism in order to regrow a fully functional limb, it will be important to study size regulation in multiple species to identify the shared mechanisms regulating this process.

## Materials and Methods

### Animal Husbandry and Surgeries

Ethical approval for this study was obtained from the Institutional Animal Care and Use Committee at the University of Massachusetts Boston (Protocol # IACUC2015004) and all experimental undertakings were conducted in accordance with the recommendations in the Guide for the Care and Use of Laboratory Animals of the National Institutes of Health. Axolotls (*Ambystoma mexicanum*) were spawned either at the University of Massachusetts Boston or the Ambystoma Genetic Stock Center at the University of Kentucky. Experiments were performed on white-strain (RRID: AGSC_101J), GFP-strain (RRID: AGSC_110J), and RFP-strain (RRID: AGSC_112J) Mexican axolotls (*Ambystoma mexicanum*). Animal sizes are measured snout to tail tip and described in the text for each experiment. They were housed in 40% Holtfreters on a 14/10-hour light/dark cycle and fed *ad libitum*. Animals were fed every day or three times a week depending on the size of the animal. Animals were anesthetized in 0.1% MS222 prior to surgery or imaging. Live images were obtained using a Zeiss Discovery V8 Stereomicroscope with an Axiocam 503 color camera and Zen software (Zeiss, Oberkochen, Germany).

To generate large and small animals, larval animals were either housed at 19°C or 4°C, which slows their growth rate. Animals were grown at these temperatures until their body lengths were approximately two-fold different, at which point the smaller animals were moved to 19°C for two weeks prior to any surgical manipulation.

### Animal Measurements

When measuring limb length and body length to determine limb proportionality and growth during regeneration, limbs were measured from the trunk/limb interception to the elbow and from the elbow to the longest digit tip (Supplemental Figure 1). Body length was measured from snout to tail tip. All measurements were taken in centimeter (cm; Supplemental Figure 1). Measurements were recorded prior to experimentation and weekly following surgical manipulation. After 5 weeks measurements were biweekly, and after 10 weeks they were taken triweekly.

### Limb Amputations and Staging of Tiny Limbs

Forelimb amputations are done mid-stylopod (mid-humerus). If the bone protruded from the amputation plane after contraction of the skin and muscle, it was trimmed back to make a flat amputation plane.

Limb regeneration stages were determined through an observation of patterning and growth rates. Limbs were considered in the “early tiny limb” stage when they reached the mid-digit stage of patterning (Iten & Bryant, 1973). Prior to this, they are considered in the “blastema” stage of limb regeneration. The transition from “early tiny limb” to “late tiny limb” is determined by a statistically significant decrease in limb length growth rate (cm/day). The regenerating limb is fully regenerated when the growth rate is no longer statistically significant from the unamputated control limbs.

### Limb Denervation Surgeries

Denervation of limbs was done by making a posterior incision on the flank, at the base of the arm, and severing and removing a piece of the three nerve bundles that come from spinal nerves 3, 4, and 5, proximal to the brachial plexus. A 2-3mm piece of the nerve bundle was cut out in an effort to delay the regeneration of the nerves into the regenerates. Partial denervations were also performed by severing 1, 2, or all three of the limb nerve bundles.

1/3 denervations were performed by severing spinal nerve 5. 2/3 denervations were performed by severing spinal nerves 4 and 5. Full denervations were performed by severing all three nerves. Mock denervations were also performed by creating the same incision on the posterior side of the limb and dissecting the nerve bundles as in a typical denervation but leaving the nerve bundles intact. Because limb denervation last approximately 7 days before nerves begin to re-innervate the limb, experimental limbs were analyzed 4 days post denervation.

### Accessory Limb Model (ALM)/ Blastema Grafting

Limbs on large (average 12 cm snout to tail tip) and small (average 6 cm snout to tail tip) GFP+ donor axolotls were amputated mid-stylopod and allowed to regenerate until mid-bud stage. At that stage, they were amputated along with approximately 2mm stump tissue, and grafted onto a regenerative permissive environment on large (12cm snout to tail tip) RFP+ host axolotls or small (6cm snout to tail tip) white host axolotls. Stump tissue was included in the graft to ensure the ectopic limb is composed primarily of donor animal cells, since stump cells contribute to the regenerate (Currie et al., 2016). The regenerative permissive environment on the large hosts was created by removing an anterior patch of full thickness skin from the stylopod and deviating a limb nerve bundle to the wound site (Endo et al., 2004, McCusker & Gardiner 2013). The blastemas on the large donor animals were substantially larger than the limbs of the small host animals, making it impossible to perform this test on the small limbs. Thus, a regenerative permissive environment on the small host axolotl was generated by removing a patch of full thickness skin from flank of the animal posterior to the forelimb. The limb nerve bundle is dissected from the limb and deviated to the flank wound site on the small animal. After the blastemas are grafted on to the host wound sites, hosts are kept on ice and misted frequently for one hour, permitting attachment of the graft.

### Neural-Modified Accessory Limb Model

We developed the neural-modified ALM assay to determine if non-neural extrinsic factors were contributing to size regulation. In the NM-ALM the nerve source and host environment are decoupled by grafting a limb Dorsal Root Ganglia (DRG) from a donor animal in lieu of the deviated nerve bundle from the host animal into the wound site, as in the standard ALM surgery. Anterior wounds are created on white strain host animals. Limb DRGs were carefully extracted postmortem from GFP+ donor animals and implanted below the proximal skin surrounding the wound site (Figure 6A). The limb axon bundle is positioned in the middle of the wound site (Figure 6A). Mid-bud staged blastemas, along with approximately 2mm of stump tissue, were amputated from white strain donor animals and immediately grafted onto the DRG nerve bundle and wound site on the host animals. Animals were kept on ice and moist for two hours following grafting to ensure attachment of the graft.

In the NM-ALM experiment reported here, we used large (average 14 cm snout to tail tip) and small (average 7 cm snout to tail tip) GFP+ and white strain siblings, generated by crossing heterozygous GFP parents, as blastema donors, DRG donors, and NM-ALM hosts. DRGs from both large and small GFP+ donor animals were dissected postmortem and grafted into anterior limb wound sites on large and small white host animals (Figure 6C). Mid-bud staged blastemas from small white strain donors were then grafted onto the NM-ALM.

### Tissue Histology and Immunofluorescence

Tissues were fixed overnight at 4°C in 4% formaldehyde (RICCA Chemical Company, Arlington, TX), decalcified in 10% EDTA (VWR, Radnor, PA) for 5-14 days depending on tissue size, rehydrated in 30% sucrose for 2 days, and embedded in Tissue-Tek OCT Compound (Sakura, Torrance, CA). The OCT blocks were then flash frozen in liquid nitrogen and stored at −20°C. They were cryo-sectioned on a Leica CM 1950 (Leica Biosystems, Buffalo Grove, IL) through the mid zeugopod at 7μm thickness. Following staining, histological stained slides were mounted using Permount Mounting Medium (Thermo Fischer Scientific, Waltham, MA). VECTASHIELD Antifade Mounting Medium (Vector Laboratories, Burlingame, CA) was used to mount coverslips on the fluorescently stained slides. Sections were imaged on the Zeiss Observer.Z1 at 20x magnification. Tile scans were taken of the entire tissue section and then stitched together using the ZenPro software (Zeiss).

### Quantification of Cell Proliferation and Cell Death

For analysis of cell proliferation in the limbs, 100ng of EdU (Roche, Basel, Switzerland) was injected into the intraperitoneal space on the flank of the animal. The limbs were then collected exactly four hours after injection and prepared for staining. Harvested tissues were process for cryo-sectioning as described above. The Roche Click-It EdU kit was used, following manufactures protocol, and co-stained with DAPI (1:1000 dilution – Sigma Aldrich, St. Louis, MO). The Roche *In Situ* cell death detection kit (Fluorescein) was used to analyze apoptosis using the manufactures’ protocol. TUNEL stained sections were also co-stained with DAPI to obtain percent cell death. Using the opensource FIJI software, the number of either proliferating (EdU+) or dying (TUNEL+) cells and DAPI+ cells were counted and used to calculate the labeling indices for each. In each section, the tissues were visually separated based on morphology into three basic categories; epidermal, skeletal (bone or cartilage), and soft tissue (all other tissue), and the labeling indices were generated for each separately. Three technical (3 sections per limb), and at least three biological replicates were performed for each sample.

### Quantification of Cell Size and ECM Size

The quantification of average cell size and extra cellular area were performed separately on the entire epidermal, muscle, and skeletal tissues in each tissue section. The epidermal and muscle tissues were identified based on morphology in Wheat Germ Agglutinin (WGA - Thermo Fischer Scientific), Rhodamine phalloidin (Thermo Fischer Scientific), and DAPI (1:1000 dilution – Sigma Aldrich) stained sections. Skeletal tissues (bone or cartilage) were identified in sections stained with Harris Hematoxylin (Sigma Aldrich), Eosin Y (Thermo Fischer Scientific), and Alcian Blue (Sigma Aldrich). To measure the average cell size of epidermis and muscle in the fluorescent images, the area within WGA plasma membrane-stained cells, where the nucleus was observed (DAPI signal), was quantified using the FIJI software and averaged (Supplemental Figure 2A). While muscle cells are elongated syncytial cells, our measurements only quantified a cross-sectional area. For the skeletal elements, the average cell area was quantified by measuring the area of Alcian blue negative areas in the element that contained a nucleus (Supplemental Figure 3A).

To analyze ECM size for the epidermis and muscle, the area of all WGA internal cell spaces (with and without DAPI) were quantified. For skeletal tissue, the area of the Alcian blue negative cell spaces were quantified. The sum of these areas was calculated to obtain the “total cellular area”. The area of the complete tissue “total tissue area” was then quantified (Supplemental Figure 3B). Since the tissue sizes can vary, percent ECM was determined through the following equation:

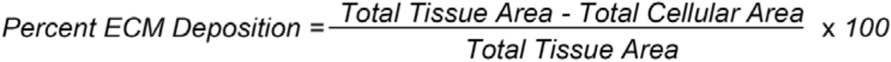

### Innervation Staining

Innervation analysis was done on regenerating limb and flank sections to quantify innervation abundance using the Mouse Monoclonal Anti-Acetylated Tubulin antibody (1:200 dilution – Sigma Aldrich), followed by the Goat-Anti-Mouse IgG Alexa Fluor 488 (1:200 dilution – Abcam, Cambridge, MA) secondary antibody. They were co-stained with Rhodamine phalloidin and DAPI as a general tissue stain, and to provide positional context for the location of the axon bundles. Limb innervation abundance was quantified by determining the percentage of limb area that is innervated (area of Anti-Acetylated Tubulin staining). To determine limb-bound axon bundle size, the sum of the area of axon bundles from DRGs 3, 4, and 5 were quantified as they emerged from the skeletal muscle surrounding the spine (Supplemental Figure 6A-B).

## Acknowledgments

The authors wish to thank Dr. Kellee Siegfried Harris, her lab members, and the members of the McCusker lab for their insightful comments during the development of this project.

## Supplemental Figures

**Supplemental Figure 1:**
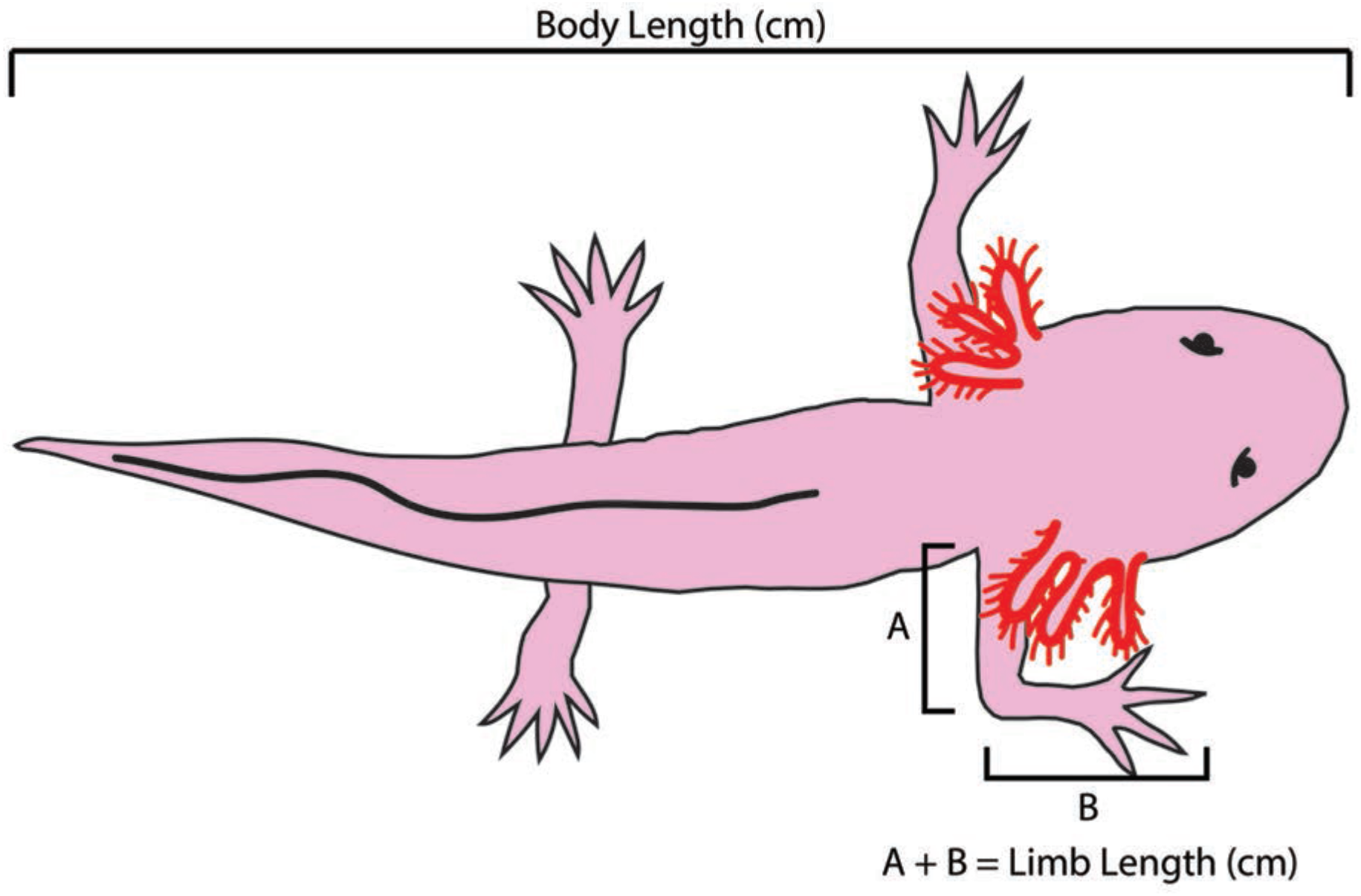
Axolotl Size measurements: Body Length is measured from snout to tail tip. Limb length is measured by measuring from the limb/trunk junction to the elbow (A) and then from the elbow to the tip of the longest digit (B). All measurements are recorded in centimeter (cm).

**Supplemental Figure 2:**
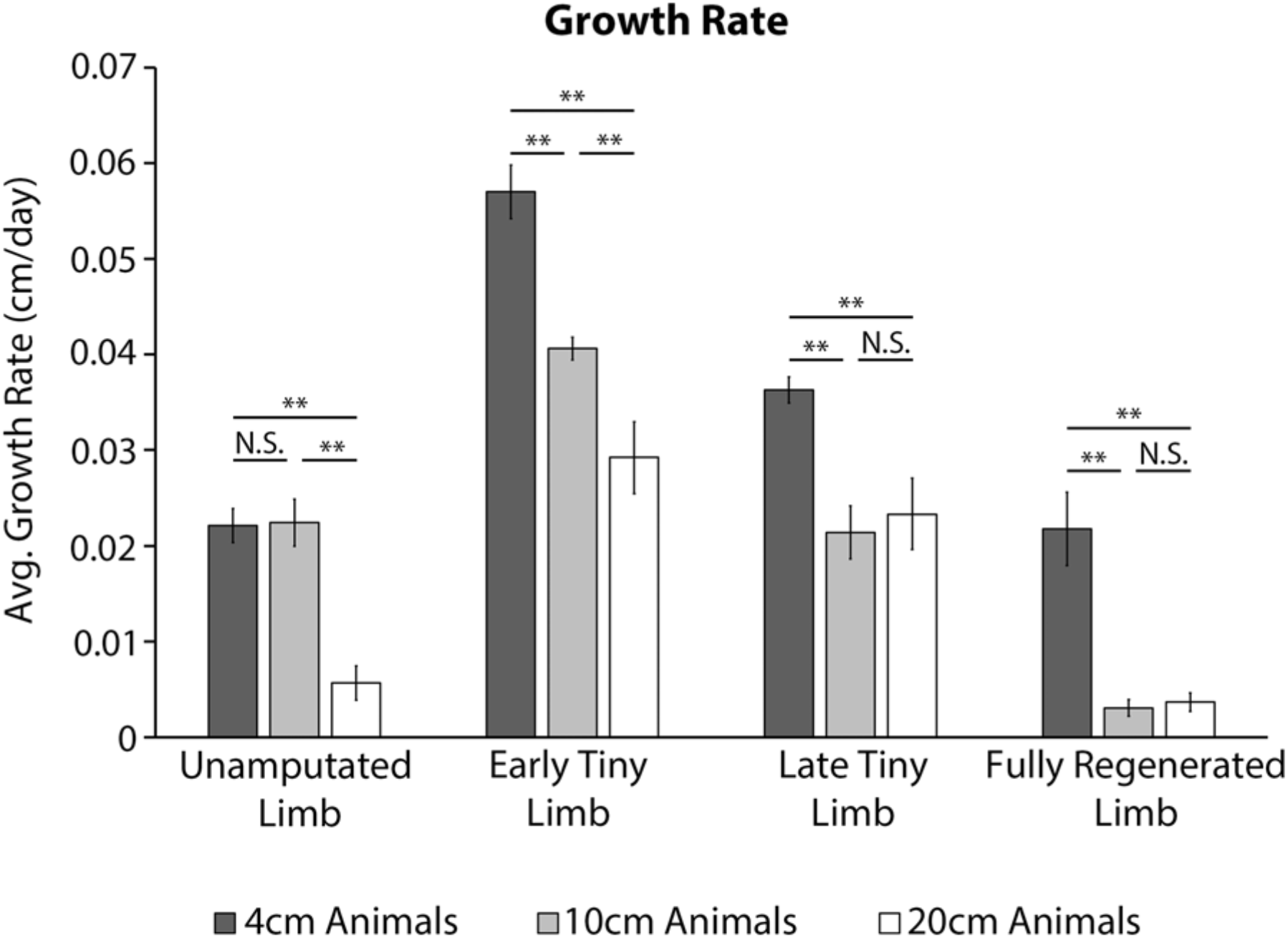
Animal size corresponds with growth rate during limb regeneration. Limb growth rates over the first 7 days of each growth phase were quantified using limb length measurements on unamputated limbs, early tiny limbs, late tiny limbs, and fully regenerated limbs of 4cm (n=20), 10cm (n=10), and 20cm animals (n=10). The body length represents their size at the time of amputation. Error bars=SEM. P-values calculated by ANOVA and the Tukey Post-hoc test. *=p<0.05 **=p<0.005.

**Supplemental Figure 3:**
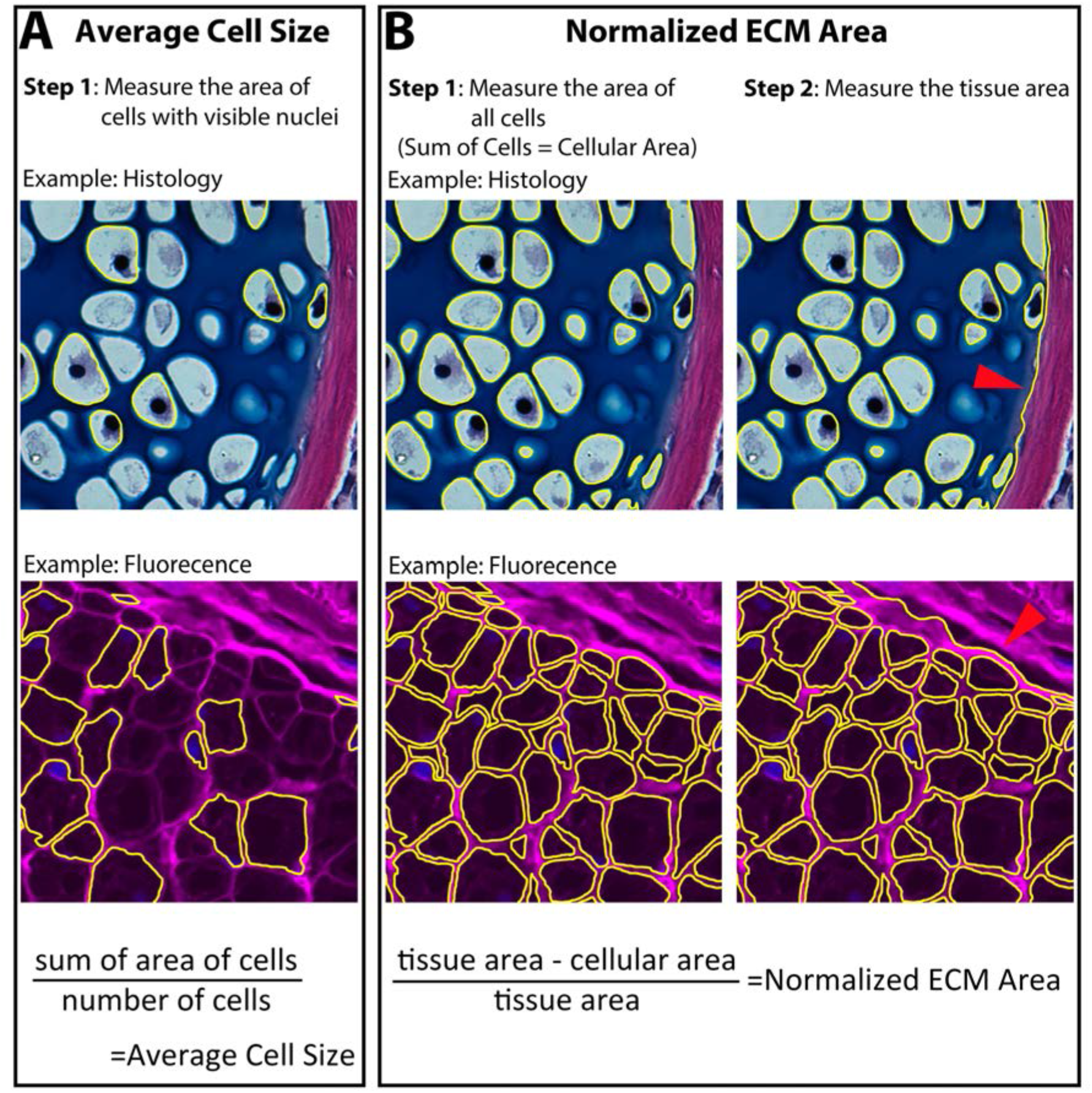
Measurement of cell and ECM size. Cell size and ECM area were quantified using 7μm cross sections through the zeugopod. Skeletal tissue was analyzed using the histology stain of hematoxylin, eosin, and Alcian blue. The epidermis and muscle (represented) were analyzed using fluorescent stains of Wheat Germ Agglutinin (WGA - magenta) and DAPI (blue). A) Average cell size was quantified by measuring the area of nucleated cells. B) Normalized ECM area was quantified by finding the sum of all the cellular area in a tissue. The sum was then subtracted from the total tissue area (red arrow indicating tissue boarder) and divided by the total tissue area to provide the normalized ECM area.

**Supplemental Figure 4:**
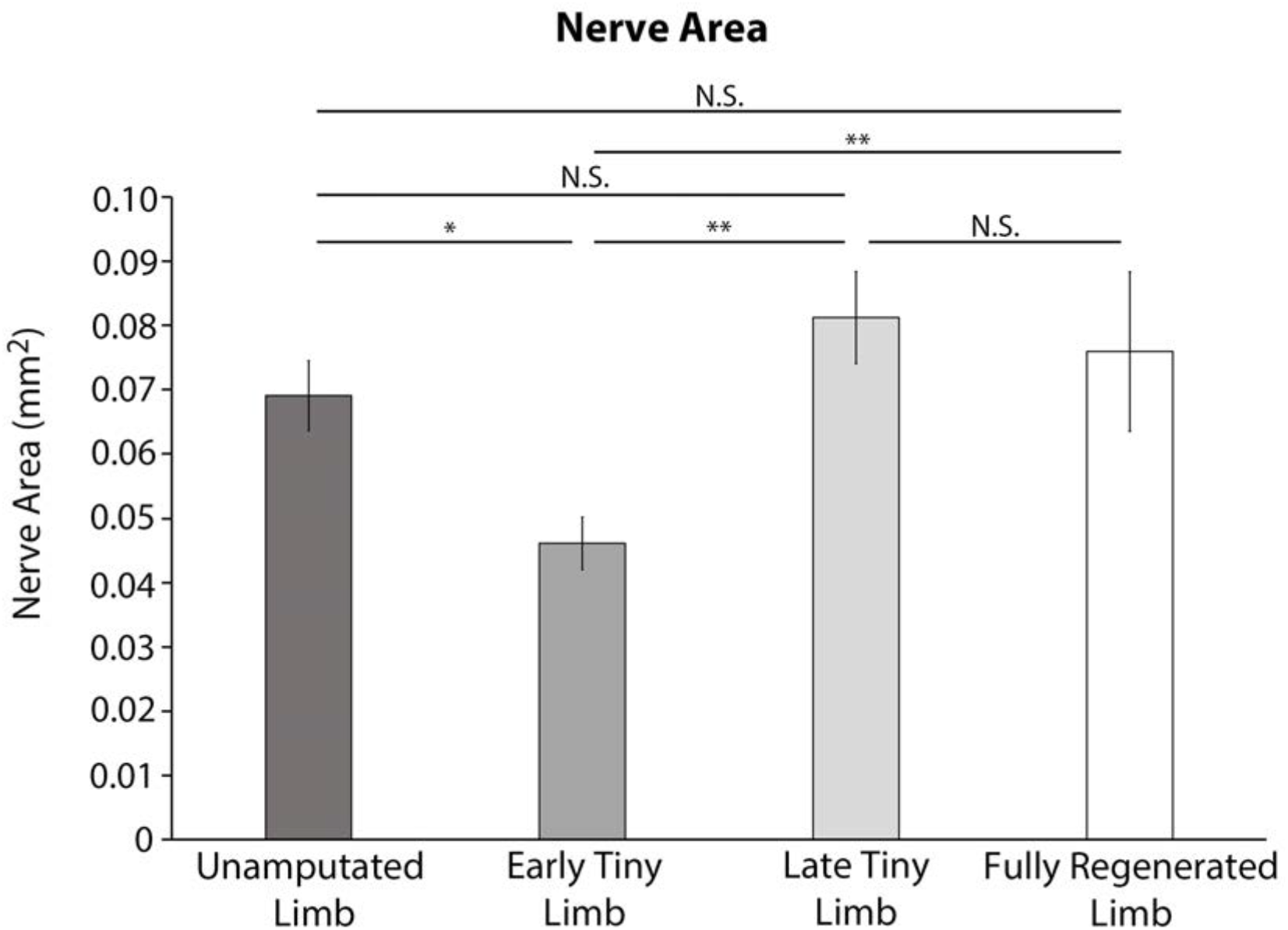
The tiny limb staged regenerate is hyperinnervated. Total nerve area was quantified from transvers limb sections of unamputated limbs, early tiny limbs, late tiny limbs, and fully regenerated limbs (n=5). Error bars = SEM. P-values calculated by ANOVA and the Tukey Post-hoc test. *=p<0.05 **=p<0.005.

**Supplemental Figure 5:**
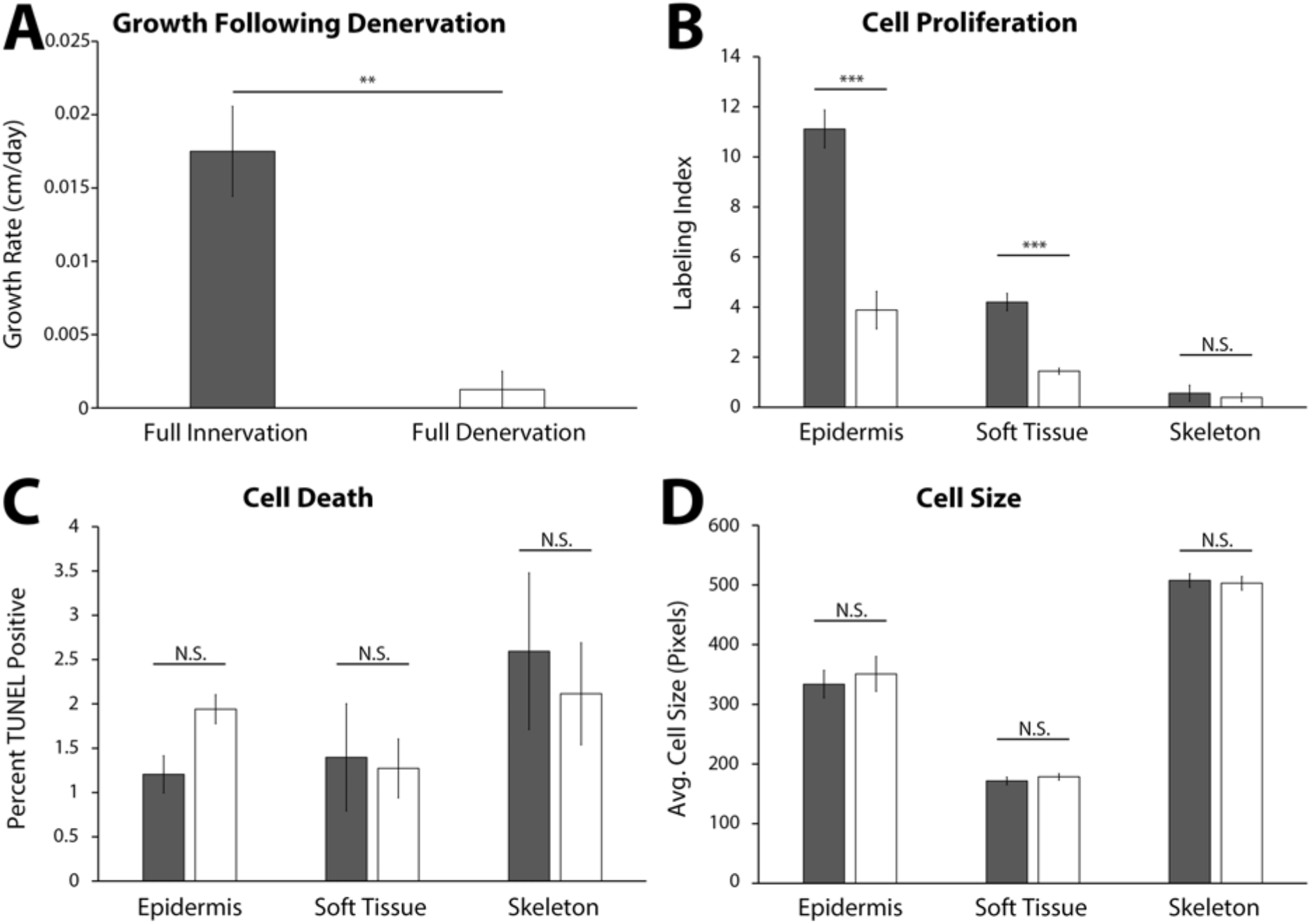
Growth of the late tiny limb requires nerve signaling for cell proliferation. Amputated limbs were permitted to regenerate to the late tiny limb stage, at which point they underwent either a mock denervation (dark grey, n=5) or a full denervation (white, n=5). The limbs were collected and analyzed 4 days post denervation for growth rate (A), cell proliferation (B), cell death (C), and cell size (D) as previously described. Error bars = SEM. P-values calculated by Paired T-Test. **=p<0.005 ***=p<0.0005.

**Supplemental Figure 6:**
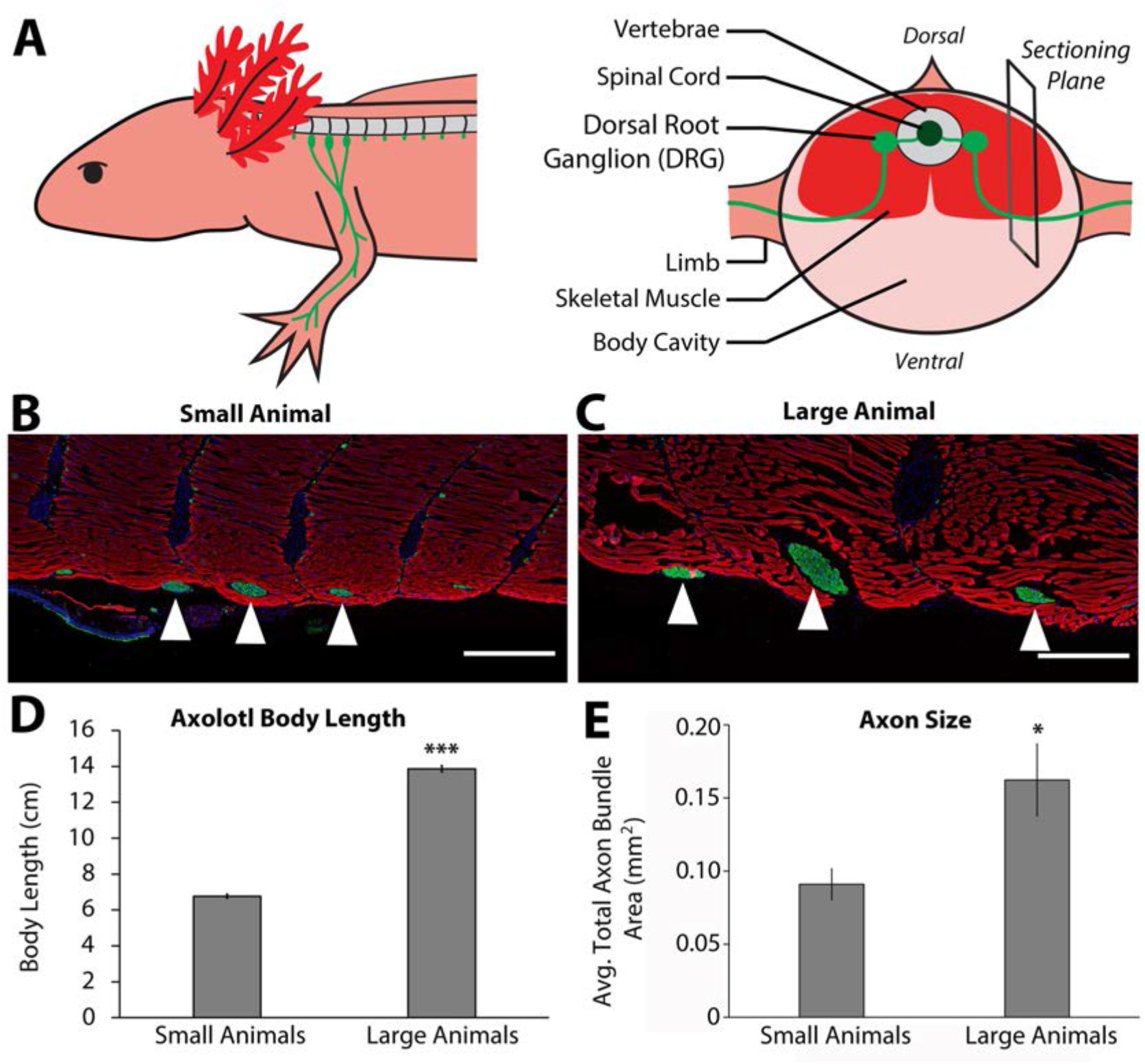
Innervation area increases with animal size. A) Three spinal DRGs (3, 4, and 5, light green) extend through the spinal skeletal muscle (red) and enter the limb. The right panel shows that the sectioning plane for B and C is located at the point where the limb-bound axon bundles emerge from the skeletal muscle. B-C) Representative immunofluorescent images of sections where the limb axon bundles (anti-acetylated tubulin antibodies – green) are emerging from the skeletal muscle (rhodamine phalloidin - red) and co stained with DAPI (blue) in both small (n=4, B) and large (n=3, C) animals. Scale bars = 500μm. White triangles indicate limb-bound axon bundles from DRGs 3, 4, and 5 (from left to right). D) Body length, snout to tail tip was measured on small (n=4) and large (n=3) animals, and the average length is represented in the graph. E) The cross-sectional area of the nerve bundles was quantified in millimeters squared, and the averages of the sum of the three axon bundles are represented in E. The large animals (n=3) have a significantly larger cross-sectional area than the small animals (n=4). Error bars=SEM. P-values calculated by T-tests. *=p<0.05 ***=p<0.0005.

**Supplemental Figure 7:**
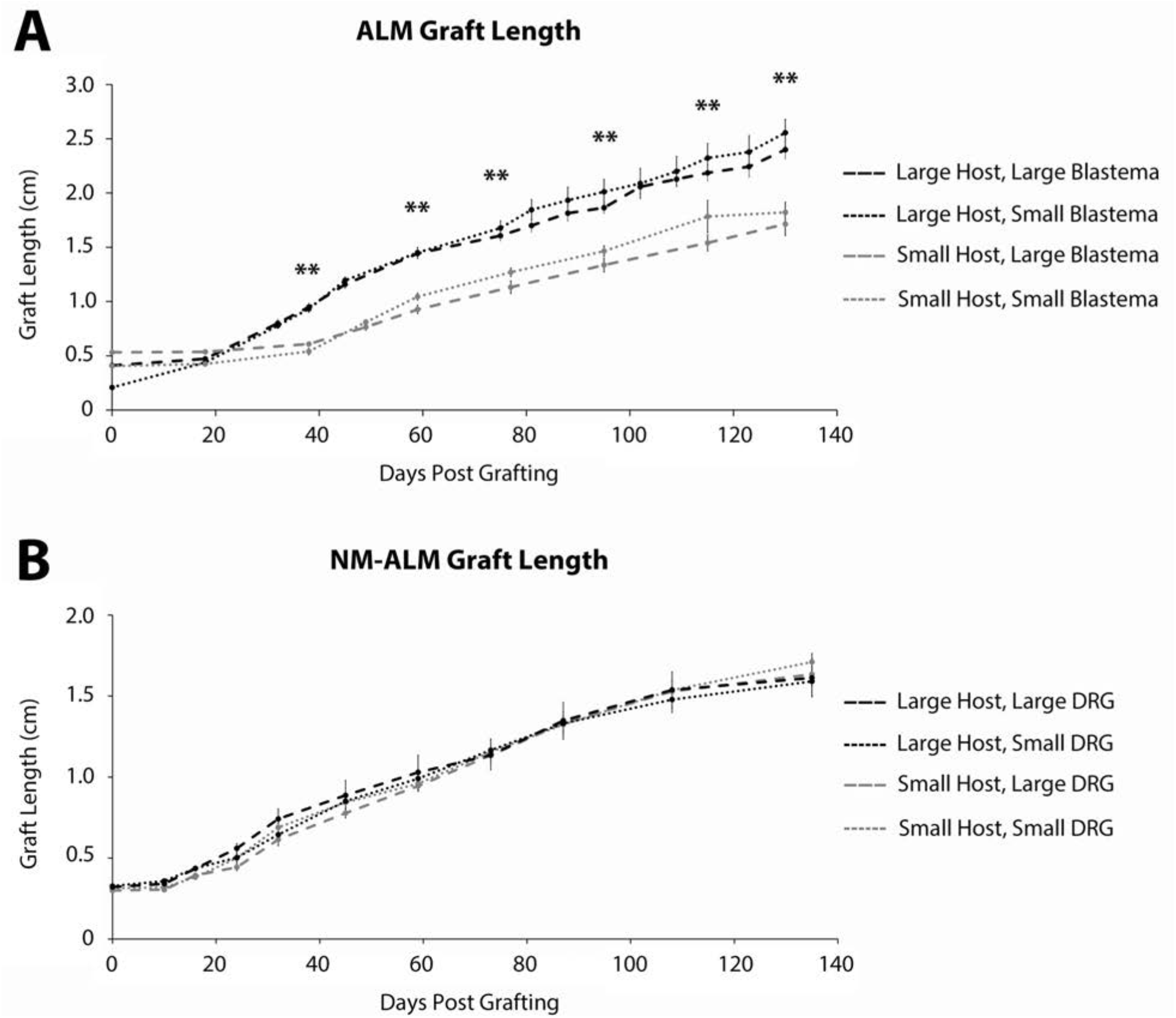
Growth of grafted limbs in ALMs and NM-ALMs. A) The ectopic limbs on the large and small host animals, generated from large and small blastema donor animals via traditional ALM surgery, were measured overtime from 0 to 130 days post grafting. B) The ectopic limbs on the large and small host animals, generated from large and small DRG donor animals via NM-ALM surgery, were measured overtime from 0 to 135 days post grafting. Error Bars = SEM. P-values calculated by ANOVA and the Tukey Post-hoc test. **=p<0.005. Statistical comparisons were made between the limbs on the differently sized host animals.

